# Cardiac Applications of CRISPR/AAV-Mediated Precise Genome Editing

**DOI:** 10.1101/2024.12.03.626493

**Authors:** Yanjiang Zheng, Joshua Mayourian, Justin S. King, Yifei Li, Vassilios J. Bezzerides, William T. Pu, Nathan J. VanDusen

## Abstract

The ability to efficiently make precise genome edits in somatic tissues will have profound implications for gene therapy and basic science. CRISPR/Cas9 mediated homology-directed repair (HDR) is one approach that is commonly used to achieve precise and efficient editing in cultured cells. Previously, we developed a platform capable of delivering CRISPR/<u>Cas</u>9 gRNAs and donor templates via <u>a</u>deno-<u>a</u>ssociated <u>v</u>irus to induce <u>HDR</u> (CASAAV-HDR). We demonstrated that CASAAV-HDR is capable of creating precise genome edits *in vivo* within mouse cardiomyocytes at the neonatal and adult stages. Here, we report several applications of CASAAV-HDR in cardiomyocytes. First, we show the utility of CASAAV-HDR for disease modeling applications by using CASAAV-HDR to create and precisely tag two pathological variants of the titin gene observed in cardiomyopathy patients. We used this approach to monitor the cellular localization of the variants, resulting in mechanistic insights into their pathological functions. Next, we utilized CASAAV-HDR to create another mutation associated with human cardiomyopathy, arginine 14 deletion (R14Del) within the N-terminus of Phospholamban (PLN). We assessed the localization of PLN-R14Del and quantified cardiomyocyte phenotypes associated with cardiomyopathy, including cell morphology, activation of PLN via phosphorylation, and calcium handling. After demonstrating CASAAV-HDR utility for disease modeling we next tested its utility for functional genomics, by targeted genomic insertion of a library of enhancers for a massively parallel reporter assay (MPRA). We show that MPRAs with genomically integrated enhancers are feasible, and can yield superior assay sensitivity compared to tests of the same enhancers in an AAV/episomal context. Collectively, our study showcases multiple applications for *in vivo* precise editing of cardiomyocyte genomes via CASAAV-HDR.

## Introduction

CRISPR-Cas9, a component of bacterial and archaeal immune systems^1^, has been engineered into a formidable molecular biology tool^2–5^, which has inspired a steady stream of advances and innovations. This success of CRISPR-Cas9 has, in large part, been due to its highly programmable DNA targeting by base pairing to guide RNAs (gRNAs)^6^. This ease of programming and high on-target editing efficiency stands in contrast to alternative gene editing approaches, such as ZFNs and TALENs. CRISPR-Cas9 has been widely adopted in diverse applications as an efficient means of creating targeted DNA double strand breaks (DSBs). Nuclease-induced DSBs are typically repaired through either non-homologous end-joining (NHEJ) or homology-directed repair (HDR). In virtually all known cell types the NHEJ pathway is favored over HDR, resulting in small, unpredictable insertions and deletions (indels) at the site of repair, which can disrupt gene function^7^. While these indels can be useful in some experimental contexts^8–11^, the creation of precise edits is often more desirable.

Precise repair of DSBs through the HDR pathway requires the presence of a homologous DNA strand, such as a sister chromatid or an exogenous donor template. Thus, by designing exogenous donor templates to induce specific edits, HDR can be used to precisely create mutations, correct mutations, or insert transgenes^12^.

Previously, we developed a platform capable of delivering CRISPR/<u>Cas</u>9 gRNAs and donor templates via <u>a</u>deno-<u>a</u>ssociated <u>v</u>irus to induce <u>HDR</u> (CASAAV-HDR). CASAAV-HDR can create precise edits *in vivo* within both mitotic and postmitotic cardiomyocytes, with neonatal editing in up to ∼45% of cardiomyocytes^13^. Here, we conduct several proof-of-concept experiments to demonstrate the broad utility of CASAAV-HDR for cardiomyocyte-specific genome editing applications. First, we used CASAAV-HDR for disease modeling by creating two different mutations in the Ttn gene, which resulted in premature stop codons. These titin truncation proteins, which have been shown to cause inherited dilated cardiomyopathy (DCM)^14,15^, were fused to the red fluorescent protein mScarlet to allow direct protein visualization. Using this novel approach we were able to rapidly observe protein localization *in vivo* at endogenous expression levels. Furthermore, in an additional disease modeling application, we used CASAAV-HDR to create an mScarlet-tagged phospholamban (PLN) variant featuring arginine 14 deletion (R14Del), which has been found to cause familial DCM^16–19^. This allowed us to assess protein localization and to characterize the effect of the mutation on cell structure, cellular function, and protein function.

In addition to disease modeling, precise genome editing can be used to functionally analyze the impact of *cis*-regulatory elements on gene expression. *Cis*-regulatory elements play dynamic roles in gene regulation during cardiac development, homeostasis, and disease. With the advent of affordable sequencing, the number of *cis*-regulatory elements that can be predicted from epigenomics data has sky-rocketed, while the ability to validate these predictions *in vivo* has historically been very limited.

To address this deficiency we pioneered the use of AAV-based massively parallel reporter assays (MPRAs) within the heart^20–22^. MPRAs allow for thousands of measurements to be made simultaneously in independent cell populations within a single sample, without need for cell sorting or complex sample processing steps. MPRAs can be deployed in several configurations, but one of the most common, termed STARR-seq, involves insertion of a library of cis-regulatory elements into the 3’ UTR of a reporter gene^23^. Each library element has a different impact on reporter gene expression, and this impact is assessed by sequencing the portion of the reporter transcript containing the element and measuring its frequency relative to other elements within the pool. Several vectors have successfully been used to deploy STARR-seq assays, including plasmids, AAV, and lentiviruses, however, all of these vectors have limitations. Plasmids are not effective for *in vivo* deployment in most tissues, AAV genomes are maintained as extra-chromosomal episomes which may not fully recapitulate regulatory effects of the native epigenomic landscape, and lentiviruses inefficiently transduce cardiomyocytes *in vivo* and integrate into the host genome in a semi-random fashion which could add locus dependent noise to the assay. Furthermore, the viral terminal repeats present in AAV and lentivirus have been shown to contain a variety of transcription factor binding sites^24–27^, and these complex elements may have transcriptional activity sufficient to alter the expression of adjacent reporter genes^27,28^. We hypothesized that we could assay the activity of precisely integrated *cis*-regulatory elements *in vivo* within cardiomyocytes in the absence of confounding effects from viral ITRs via CASAAV-HDR mediated insertion of a library of regulatory elements into the 3’ UTR of a native gene, in the fashion of a STARR-seq experiment. In addition to demonstrating the utility of CASAAV-HDR for disease modeling, here we present data confirming the feasibility of this MPRA strategy.

## Results

### CASAAV-HDR for disease modeling: Titin truncation mutants

Pathogenic variants in the *TTN* gene, which encodes the massive sarcomere protein titin, have been under-analyzed due to the size limitations of common vectors. TTN-truncating variants (TTNtvs) are the most common genetic cause of DCM, occurring in ∼25% of familial cases and ∼18% of sporadic cases^14,29^. However, the mechanism by which TTNtvs cause DCM remains controversial^30^. Various hypotheses have been proposed, with some studies indicating that TTNtvs cause DCM through a dominant negative mechanism, by which truncated titin is expressed and incorporated into the sarcomere^31,32^; while others suggest a mechanism of haploinsufficiency, whereby premature stop codons trigger nonsense-mediated decay, which leads to insufficient titin protein generated^14,33^.

Therefore, determining whether truncated titin proteins are present and documenting their localization is paramount to distinguishing the two mechanisms and improving our understanding of how TTNtvs cause DCM.

Previously we demonstrated that CASAAV-HDR could be used to fuse the red fluorescent protein mScarlet to the wildtype titin C-terminus (WT TTN)^13^. Here we used CASAAV-HDR to study two different TTNtvs, T deletion (TTN-TDel) and AT insertion (TTN-ATIn) in Exon 326 (mouse exon 274), both of which result in a frameshift, and have been reported to associate with DCM^14,15,29^. This was achieved by engineering AAV9 HDR vectors containing a sgRNA and a donor template that inserts mScarlet upstream of the premature or native stop codons (Figure 1A). Thus, upon successful HDR a mutant TTN-mScarlet fusion protein is produced, allowing for subcellular protein localization to be analyzed.

**Fig. 1.**
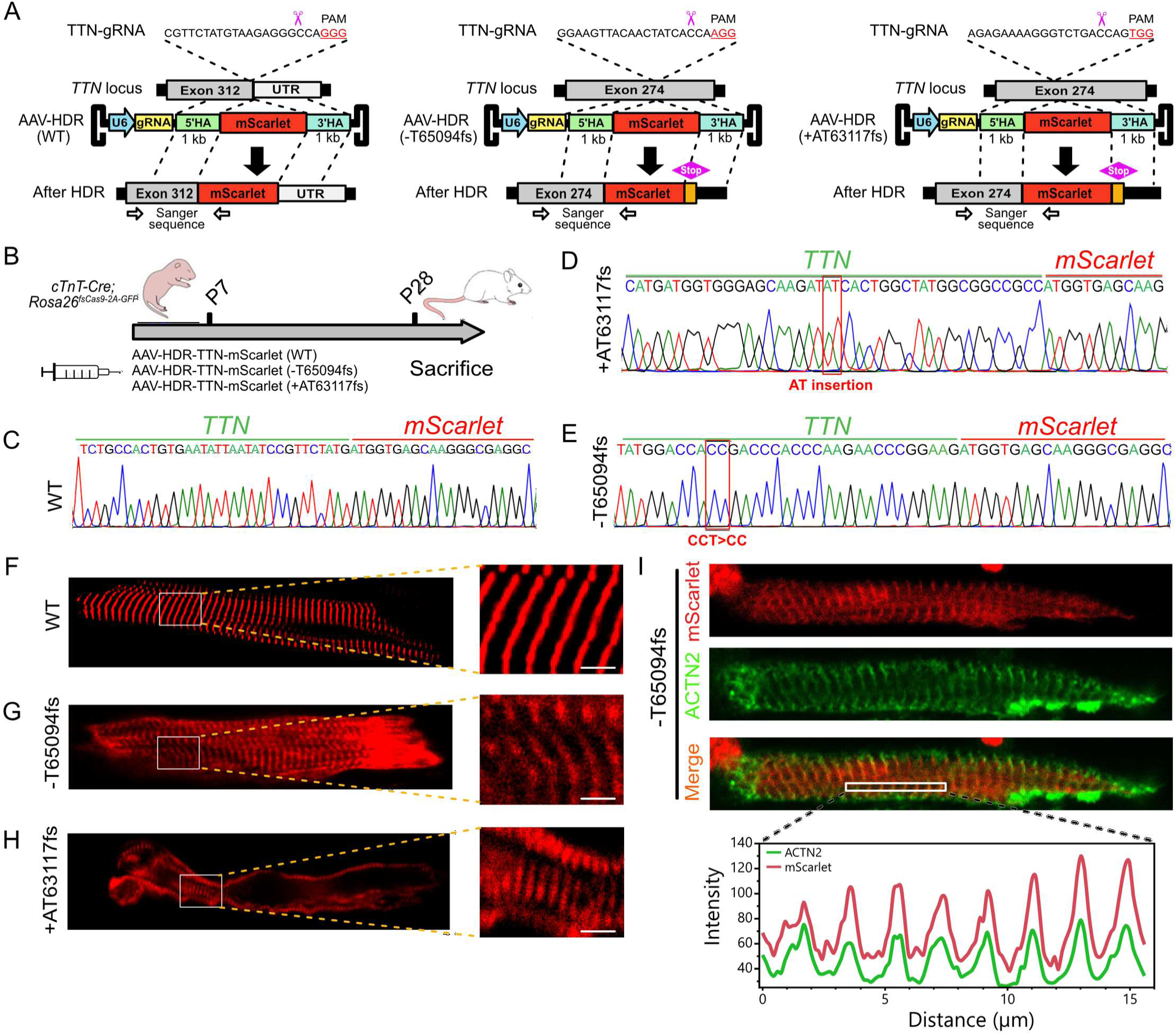
Creation of titin truncating variants via CASAAV-HDR. **A.** Genome editing strategy for creation of titin variants or control alleles, with c-terminal insertion of mScarlet tag. **B.** Experimental timeline. **C-E.** Exon-mScarlet junction sequences from wildtype control, +AT63117fs, or-T65094fs cDNA, respectively. **F-H.** In situ imaging of mScarlet fluorescence in cardiomyocytes with Ttn wildtype control,-T65094fs, or +AT63117fs alleles, respectively. Scale bar = 2.5μm. **I.** Isolated T65094fs cardiomyocyte immunostained with alpha actinin (ACTN2). ACTN2 signal colocalizes with mScarlet (lower panel).

AAV9 was subcutaneously injected into mouse pups with cardiac-restricted Cas9 expression (*Tnnt2-Cre; Rosa26^fsCas9-P2A-GFP/+^*) at postnatal day 7, and mScarlet expression was analyzed at postnatal day 28 (Figure 1B). First we confirmed that CASAAV-HDR succeeded in precisely inserting mScarlet into the target loci, creating the desired truncation variants. The 5’ junctions between the inserted mScarlet and the endogenous TTNtvs or TTN-WT were amplified from cardiomyocyte cDNA and Sanger sequenced. Sanger sequencing showed successful precise insertion of mScarlet at the target loci for all three alleles (Figure 1C-E). As previously reported^13^, *in situ* confocal imaging revealed that ∼30% of ventricular TTN-WT cardiomyocytes strongly expressed TTN-mScarlet, which robustly integrated into the sarcomere (Figure 1F). Next we analyzed hearts injected with the TTNtvs vectors, and in both cases the truncated TTN-mScarlet fusion protein was detected. mScarlet localized at least in part to the sarcomere, suggesting that the variants are stable and sarcomerically integrated (Figure 1G-H).

Titin spans half of a sarcomere, with the N-terminus aligned to the Z-line and the C-terminus at the M-band^34,35^. Truncated TTN-TDel is approximately two-thirds the length of wildtype TTN; therefore, we expected mScarlet fluorescence to localize between the Z-line and M-line. Surprisingly, we found that fluorescence from TTN-TDel-mScarlet closely co-localized with α-Actinin (Figure 1I) in isolated immunostained cardiomyocytes, suggesting that the titin N-terminus incorporated into the Z-line, while the fluorescently labeled C-terminus fails to extend as it normally should towards the M-band.

Collectively, these data demonstrate that CASAAV-HDR can be used to quickly and precisely create and label disease relevant mutant gene products with fluorescent reporters, which can be used to monitor protein level and localization. We observed reporter-labeling of both WT and two mutant titin variants, and found that the TTNtvs could be stably expressed and at least partially integrated into the sarcomere, supporting a potential dominant-negative DCM disease mechanism for TTNtvs.

### CASAAV-HDR for disease modeling: Creation of a phospholamban point mutation

Phospholamban (PLN), a small, reversibly phosphorylated transmembrane protein localized in the sarcoplasmic reticulum membrane (SR), is a crucial regulator of the activity of the calcium pump sarco(endo)plasmic reticulum Ca^2+^-ATPase type 2a (SERCA2a)^36^. SERCA2a clearance of Ca^2+^ from cytosol into the SR is inhibited by unphosphorylated PLN. PLN phosphorylation relieves this inhibition, resulting in enhanced Ca^2+^ reuptake during diastole and facilitating cardiac relaxation.

Deletion of arginine 14 (R14Del) impairs PLN phosphorylation and causes DCM^16–19^. The optimal approach for in vivo studies of such a mutation is to knock the mutation into the endogenous mouse gene. However, creating knockin mice is time consuming and costly. The mutant protein could also be overexpressed using transgenic or AAV approaches. However, these strategies suffer from potential overexpression artifacts. We hypothesized that CASAAV-HDR could be used to overcome these limitations by enabling rapid and cost-efficient creation of gene-targeted mutant alleles.

As a proof-of-concept demonstration of CASAAV-HDR’s value for disease modeling, we engineered a vector containing a PLN 5’ UTR-targeting sgRNA and a mutant (PLN-R14Del) or wildtype control (WT) promoterless donor template. Upon successful HDR, mScarlet and the PLN WT or R14Del sequence is inserted into the N-terminus of the native PLN coding sequence (Figure 2A).

**Fig. 2.**
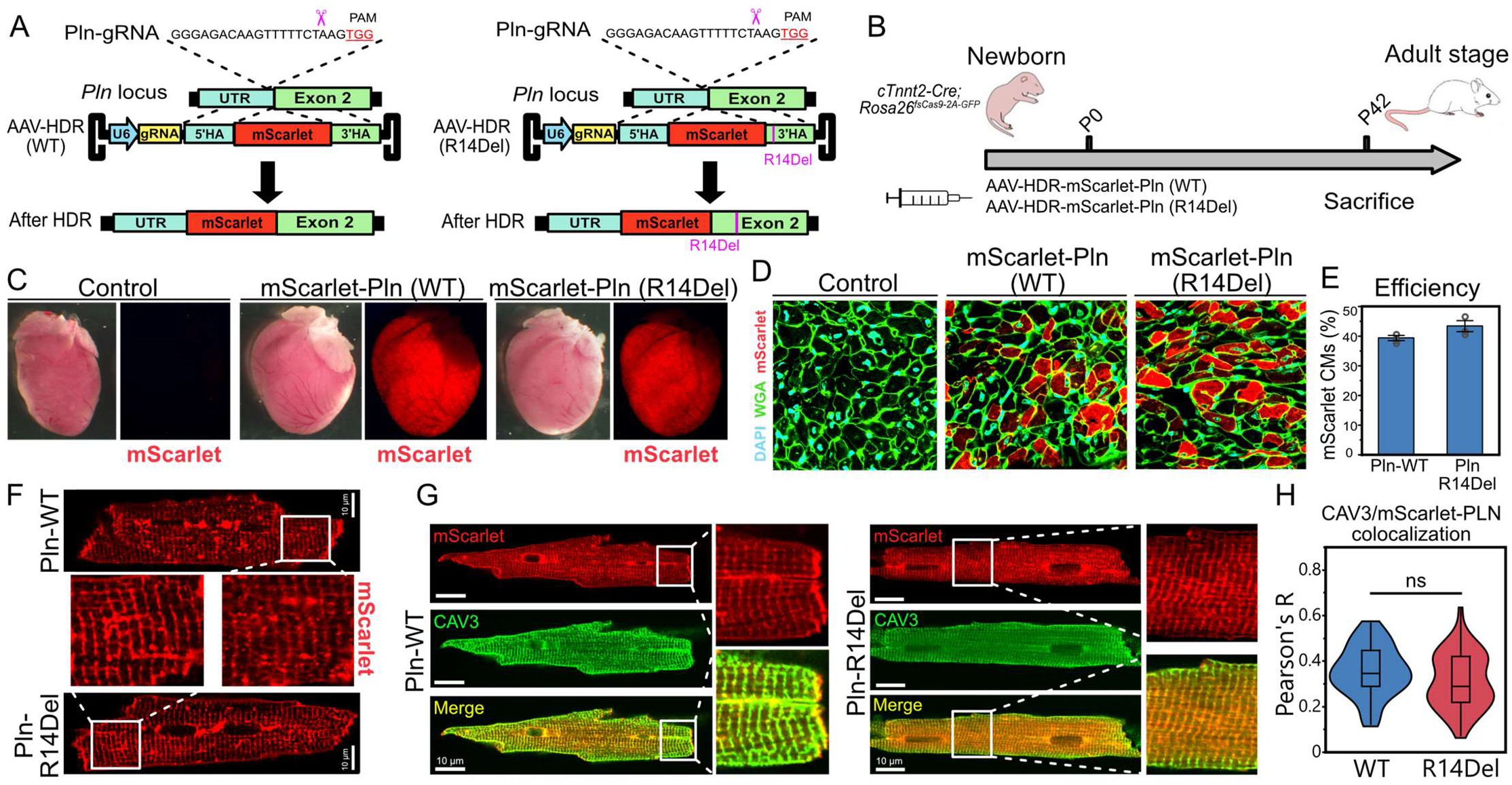
Creation of a phospholamban point mutation via CASAAV-HDR. **A.** Homology directed repair-based strategy for genome editing at the Pln locus. **B.** Experiment timecourse. Guide RNA and donor template was delivered by AAV9 to newborn mouse pups with cardiomyocyte specific Cas9 activation. **C.** Wholemount images of 6-week old uninjected control or Pln edited hearts. **D.** DAPI and WGA stained ventricular sections of control and edited hearts. **E.** Editing efficiency as the percent of ventricular cardiomyocytes that expressed mScarlet. **F.** In situ confocal imaging of edited cardiomyocytes. **G.** mScarlet-expressing cardiomyocytes immunostained with t-tubule marker CAV3. **H.** Colocalization analysis of CAV3 and mScarlet-PLN in immunostained cardiomyocytes. Student’s t *p =* 0.052. n = 60, and 55 for WT and R14Del groups, respectively.

This AAV9-based HDR vector was delivered by systemic injection to cardiomyocytes of newborn *Tnnt2-Cre; Rosa26^fsCas9-P2A-GFP^* mice, and at six weeks of age, hearts were collected (Figure 2B). Analysis revealed robust cardiac mScarlet expression in both WT and R14Del groups (Figure 2C). Quantification of knockin efficiency from ventricular sections revealed that ∼39% and ∼43% of cardiomyocytes were mScarlet+ in PLN-WT and PLN-R14Del hearts, respectively (Figure 2D,E). The intensity of mScarlet fluorescence in PLN-WT hearts was elevated in the ventricle relative to the atria (Supp. Figure 1A), consistent with the known expression profile of PLN (Supp. Figure 1B)^22^. Heart sections from Cas9 negative mice injected with the CASAAV-HDR PLN-WT vector displayed negligible mScarlet fluorescence, indicating that fluorescence in Cas9 positive groups does not originate from transcription of the AAV vector (Supp. Figure 1C). Amplification and Sanger sequencing of ventricular cDNA verified the programmed junction between the mScarlet insertion and the downstream native exon 2 sequence (Supp. Figure 1D-F).

Previous studies in cell lines and transgenic mice have reported a variety of localization profiles for PLN-R14Del, including aberrant cytoplasmic localization^17^, aberrant localization to the plasma membrane^18^, or no effect on localization^37^. In gene targeted PLN^R14Del/+^ mice, no difference in localization was observed at 8 weeks of age, while moderate levels of PLN aggregation were observed by 20 months^38^. In contrast, PLN^R14Del/R14Del^ mice exhibited a much stronger phenotype, including dramatic PLN aggregation by 8 weeks of age. To assess protein localization with our approach, we employed *in-situ* confocal imaging of mScarlet-labeled cardiomyocytes, and observed similar subcellular localization profiles for PLN-R14Del and PLN-WT proteins. In both PLN-WT and PLN-R14Del cardiomyocytes, mScarlet-PLN localized to the sarcoplasmic reticulum, with minor protein aggregation observed (Figure 2F). To more quantitatively characterize PLN localization, we isolated the cardiomyocytes from PLN-R14Del and PLN-WT hearts and immunostained them with Caveolin-3 (CAV3), a protein that localizes to the T-tubules, in tight proximity to the junctional sarcoplasmic reticulum. mScarlet-PLN colocalization with CAV3 was comparable between PLN-WT and PLN-R14Del cardiomyocytes (Figure 2G,H), indicating that the localization of mScarlet-PLN is not disturbed by the R14Del mutation in CASAAV-HDR edited cardiomyocytes at 6 weeks of age.

Given the reported phenotypes from gene-targeted PLN-R14Del mice, which did not show localization disruption in heterozygous cardiomyocytes at eight weeks^38^, our results are consistent with expectations that most CASAAV-HDR PLN R14Del cardiomyocytes are heterozygous for the edit.

We next assessed phenotypes of PLN-R14Del hearts. Differences in gross morphology or heart size between PLN-WT and PLN-R14Del hearts were not observed, likely due to only a minority of cardiomyocytes being edited. However, analysis of isolated cardiomyocyte morphology revealed that the PLN-R14Del mutation slightly increased cell area, due to an increase in width (Figure 3A,B) but not length (Supp. Fig. 1G). Next we assessed t-tubule organization, which is easily disrupted by a variety of pathological insults, including hypertrophy^39^. Computational analysis of T-tubule organization using Auto-TT^40^ showed that T-tubule network regularity and transverse element spacing were unchanged in R14Del cardiomyocytes (Figure. 3C,D). An alternate metric of network organization, Auto-TT integrity index, was also unchanged (data not shown), collectively indicating that, at six weeks post-editing, R14Del cardiomyocytes have preserved T-tubule organization.

**Fig. 3.**
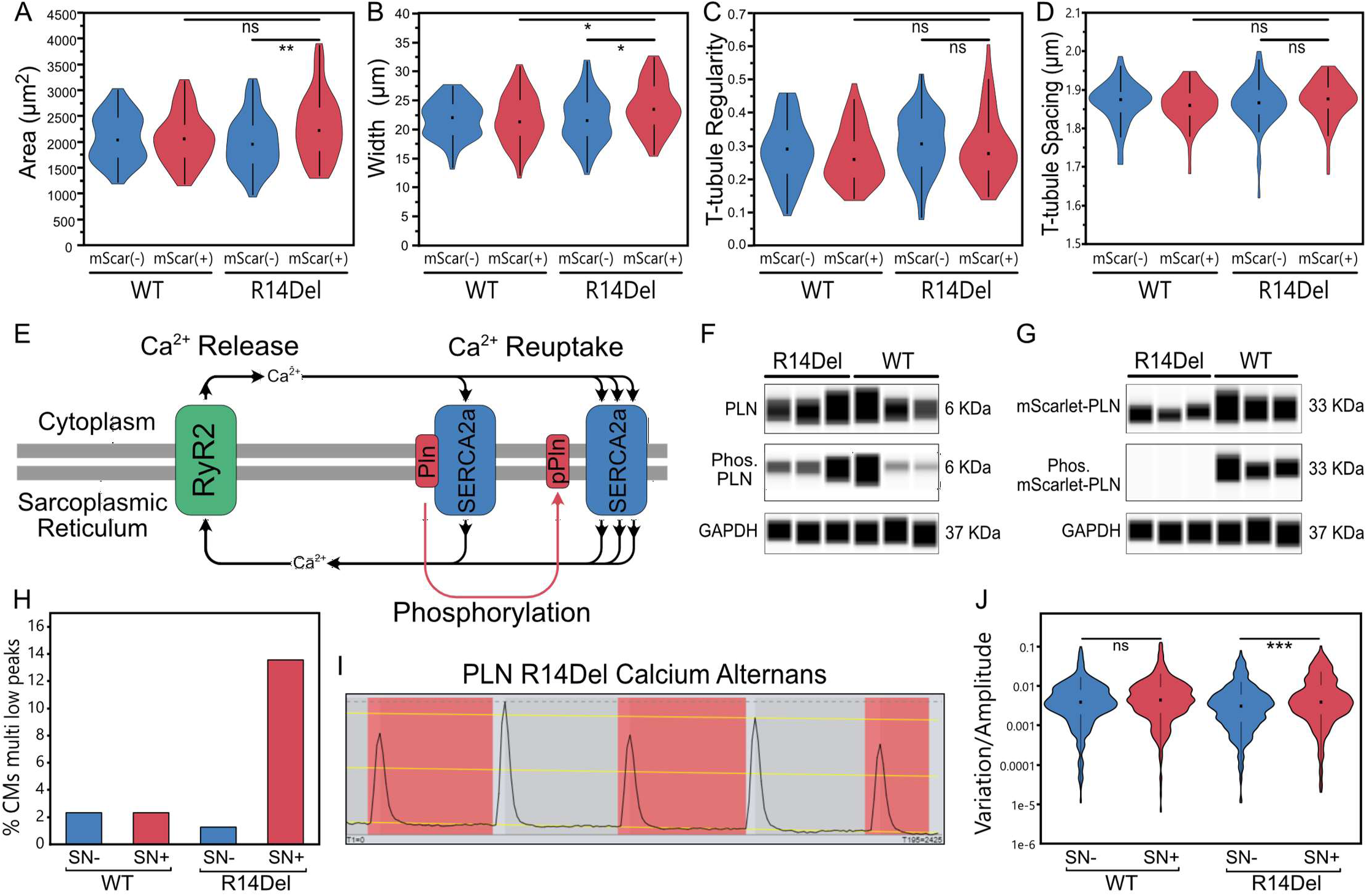
Phenotypes of PLN R14Del cardiomyocytes. **A.** Area of edited (mScarlet+) and unedited (mScarlet-) cardiomyocytes. **B.** Width of cardiomyocytes. **C.** Organization of the cardiomyocyte t-tubule network. **D.** Spacing of transverse t-tubule elements. **E.** Schematic of molecular regulation of calcium release and reuptake. **F.** Capillary western blot showing that unedited PLN is phosphorylated. **G.** Capillary western blot showing that R14Del blocks phosphorylation of PLN. **H.** Quantification of cardiomyocytes with three or more low peaks. **I.** AnomalyExplorer; beat-to-beat variation in calcium amplitude for a representative SNAP-tag(+) PLN-R14Del cardiomyocyte with multiple low peaks (highlighted in red). **J.** Quantification of variation in peak height. For each set of calcium traces peak coordinates were fit with a best-fit line. The distance of each peak from the best fit line was normalized to peak height and plotted. n = 429, 430, 395, and 295, for WT SN-, WT SN+, R14Del SN-, and R14Del SN+, respectively. p<0.05*; <0.01**; <0.001***; not significant, ns. A-D) Dunnett’s p-value multiple comparison to R14Del mScarlet(+) group. n = 56, 63, 63, and 62, for WT SN-, WT SN+, R14Del SN-, and R14Del SN+, respectively. J) One-way ANOVA, Tukey multiple comparison correction.

We next evaluated mechanisms and functional phenotypes in PLN-R14Del cardiomyocytes, first by assessing a key regulatory mechanism of PLN activity: phosphorylation at Ser16 and Thr17 (Figure 3E)^41^. Capillary western blot analysis of ventricular homogenates from PLN-WT and PLN-R14Del mice showed that a subset of unedited PLN (6 KDa) was phosphorylated (pPLN; 6 KDa) in both groups (Figure 3F). In contrast, mScarlet-PLN (33 KDa) was only phosphorylated in the WT group (Figure 3G), demonstrating that, consistent with past reports, R14Del blocks PLN phosphorylation at Ser16 and Thr17^37,42^.

Given the crucial nature of PLN phosphorylation in cardiomyocyte calcium cycling, we next examined calcium handling in PLN-WT and PLN-R14Del cardiomyocytes. To enable analysis of a calcium indicator in the red channel, we first designed CASAAV-HDR donor templates that replaced mScarlet in our original iteration with a SNAP-tag (Supp. Figure 2A). As before, SNAP-WT and SNAP-R14Del vectors were injected at birth and isolated cardiomyocytes were analyzed at six weeks of age (Supp. Figure 2B). Isolation and Sanger sequencing of cDNA from edited cardiomyocytes confirmed precise insertion of the SNAP-tag and creation of the R14Del mutation (Supp. Figure 2C,D). Incubation of SNAP-PLN edited cardiomyocytes with a cell-permeable far-red SNAP-tag-binding dye efficiently marked live edited cells (Supp. Figure 2E), allowing for measurement of calcium transients in the red channel using Rhod-2AM (Supp. Figure 2F,G). First we quantified standard Ca^2+^ transient metrics including peak amplitude, upstroke velocity, downstroke velocity, and action potential duration at 50% of the peak. No significant differences between SNAP-PLN-WT and SNAP-PLN-R14Del cardiomyocytes were observed (Supp. Figure 2H-K). Since humans with the R14Del mutation exhibit elevated susceptibility to arrhythmias, we next analyzed our data set for the occurrence of abnormal Ca^2+^ events, which are commonly observed in arrhythmogenic phenotypes^43^. Exploration of our dataset using the analysis package AnomalyExplorer^44^ indicated that SNAP-PLN-R14Del cardiomyocytes had increased variation in peak amplitude, evidenced by an increased percentage of cells with multiple “low peaks” relative to control groups (Figure 3H). Visual analysis of traces with multiple low peaks revealed that nearly all had a pattern of beat-to-beat alternation between higher and lower peaks (Figure 3I), consistent with descriptions of Ca^2+^ alternans, a highly arrhythmogenic phenomenon linked to SERCA inhibition^45–47^. To better quantify this phenotype, we directly measured the variation in peak amplitude across all sample groups and found that SNAP-PLN-R14Del cells had significantly higher variation in amplitude than unedited cells from the same hearts (Figure 3J; *p*=0.0007). In comparison, no difference was observed between edited (SNAP+) and unedited (SNAP-) PLN-WT cells (*p*=0.1514).

These data show that the PLN-R14Del mutation can be efficiently generated *in vivo* within postnatal cardiomyocytes via CASAAV-HDR, allowing for rapid and robust localization analysis. We also show that by six weeks after creation of the mutation, structural defects are mild, while PLN phosphorylation showed marked impairment. This impairment likely contributed to the observed Ca^2+^ alternans phenotype.

### CASAAV-HDR for precisely integrated massively parallel reporter assays

Having demonstrated the utility of CASAAV-HDR for disease modeling, we next assessed the technique’s potential to enable a previously unattainable functional genomics assay: an MPRA featuring candidate enhancers that have been precisely integrated into genomic DNA. To execute this unique MPRA, we used AAV-mediated HDR to precisely integrate a pool of enhancers into the 3’ UTR of the mouse *Tnni1* gene, in the fashion of a STARR-seq experiment. *Tnni1* is highly expressed in immature cardiomyocytes, but is nearly silent in adult cardiomyocytes (Sup. Fig. 3A)^21^. We anticipated that insertion of an active enhancer into the *Tnni1* UTR would result in persistently elevated expression of *Tnni1* in adult cardiomyocytes. We designed an enhancer pool consisting of 400 bp regions from 25 validated cardiomyocyte enhancers, 25 negative control endocardial enhancers, and 5 negative control regions strongly bound by P300 in embryonic stem cells (ESC), all selected from the VISTA Enhancer Database or the literature (Supplemental Table 4)^48–59^. We analyzed P300 occupancy of the selected regions based on previously generated cardiomyocyte-specific and endothelial-specific ChIP-sequencing datasets^21,49^. As expected, P300 was selectively bound at cardiomyocyte and endothelial cell enhancers in cardiomyocytes and endothelial cells, respectively, and no P300 enrichment was observed at ESC enhancers (Fig. 4a). This pool of enhancers was synthesized with 3’ barcodes and cloned into a CASAAV-HDR donor vector targeting the *Tnni1* 3’ UTR (Fig. 4b). In this system, HDR mediates replacement of the *Tnni1* stop codon with an HA-tag, stop codon, enhancer candidate, and barcode. This donor template pool was packaged into AAV9 and delivered to *Tnnt2-Cre; R26^fsCas9-2A-GFP^* newborn mice, in which *Tnnt2-Cre* activates Cas9 expression selectively in cardiomyocytes. At one month of age, cardiomyocytes were isolated in duplicate, and a portion of the cells were fixed and immunostained with HA-tag antibody. We observed weak signal in a small fraction of cardiomyocytes (Sup. Fig. 3B; <1%), suggesting that enhancer insertion resulted in persistent *Tnni1* expression in at least a subset. From the remaining cardiomyocytes, the barcoded amplicon was amplified from genomic DNA, and RNA was reverse-transcribed and sequenced. Sequence analysis showed that the frequencies of barcoded enhancers within replicate samples were well correlated (r^2^>0.8), indicating that the enhancer pool was adequately sampled (Sup. Fig. 3C). To calculate enhancer activity, the frequency of each enhancer in RNA was normalized to its frequency in genomic DNA. We observed that cardiomyocyte enhancers showed higher activity within cardiomyocytes than negative control enhancers (Fig. 4C; on average, ∼4.5 fold higher than endocardial enhancers). Approximately half of the cardiomyocyte enhancers had an RNA:DNA ratio > 1, while only a single endocardial enhancer, and no ESC enhancers, reached this activity level. The observation that only a subset cardiomyocyte enhancers had high activity is likely explained by the tested enhancers being only 400 bp segments of the full-length validated enhancers, which averaged ∼1.99 kb, and most transgenic enhancer validation was performed in embryonic rather than the adult heart.

**Fig. 4.**
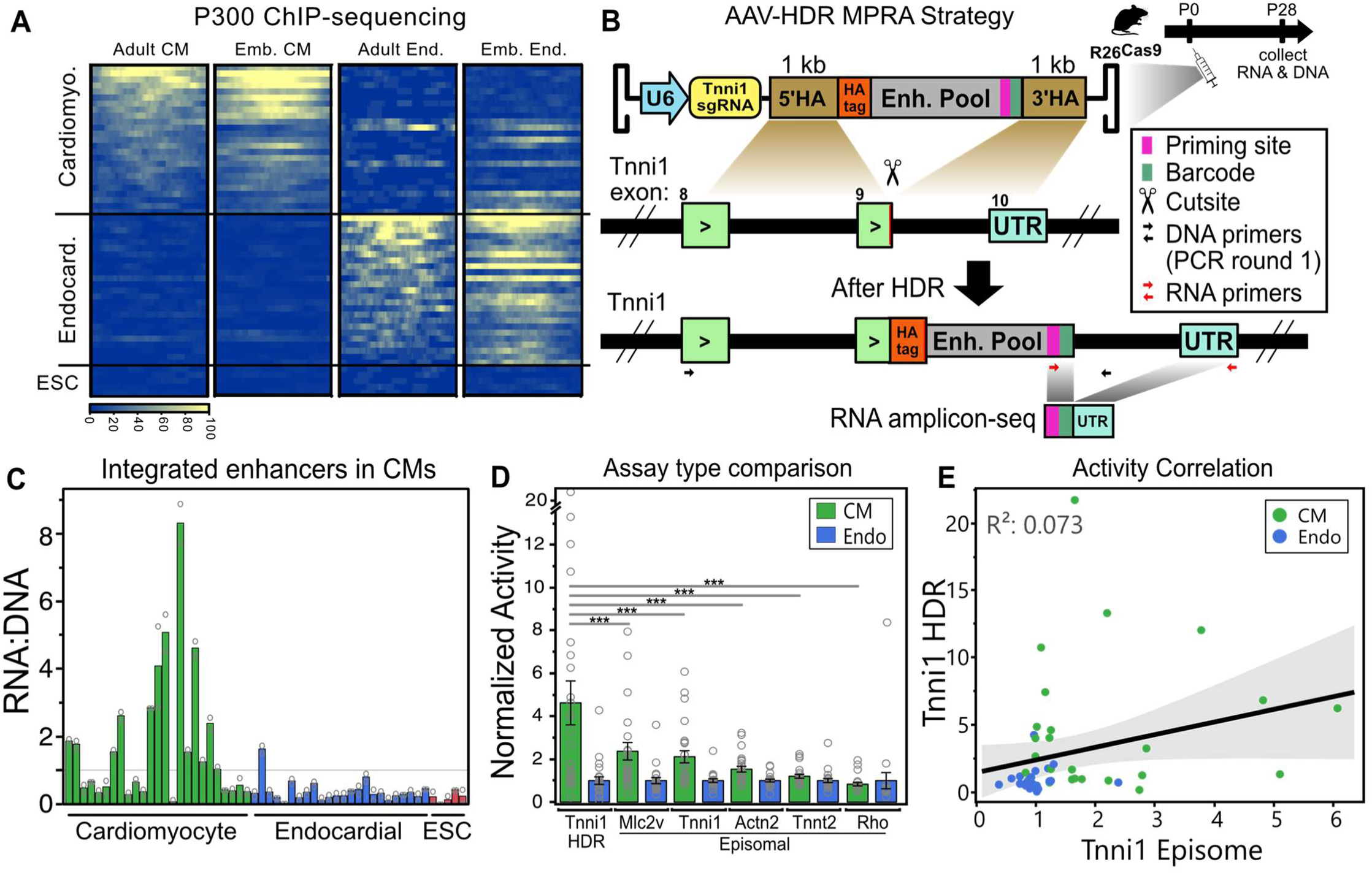
Precise integration of enhancers for a massively parallel reporter assay. **A.** P300 binding intensity at each selected enhancer in either cardiomyocytes or endothelial cells. **B.** HDR/MPRA strategy. To avoid amplification of AAV-derived DNA or transcripts, at least one primer in each set targets a region outside the homology arms. **C.** The activity of enhancers inserted into the Tnni1 3’ UTR within cardiomyocytes. **D.** Comparison of enhancer activities when inserted into the Tnni1 UTR, or when assayed in an episomal AAV STARR-seq vector with an assortment of minimal promoters. Dunnett’s p-value multiple comparison of cardiomyocyte enhancers for each promoter to the cardiomyocyte enhancers of the Tnni1 HDR MPRA. \*\*\**p*<0.001 **E**. The activities of enhancers assayed by Tnni1 HDR MPRA poorly correlate with activities obtained by episomal MPRA. Shaded area indicates confidence of fit for best-fit line.

We next sought to determine if our integrated MPRA results differed meaningfully from AAV episome-based assays. To answer this question, the enhancer pool was cloned into an AAV-vector in combination with several different minimal promoters (Supp. Table 5) and an mCherry reporter gene (Supp. Fig. 3D). The MPRA pools were packaged into AAV9, injected into wildtype mice at birth, and ventricles were collected at one month of age. After amplicon-sequencing the barcoded region of reporter gene transcripts, we calculated the activity of each enhancer with each minimal promoter, and observed that similar to the *Tnni1*-integrated MPRA dataset, the cardiomyocyte enhancers demonstrated increased activity relative to control enhancers (Fig. 4D). Of all promoters tested, the Mlc2v promoter showed the largest dynamic range, with ∼2.5 fold higher average activity in positive control cardiomyocyte enhancers than in negative control endothelial enhancers. This is significantly lower than the 4.5 fold observed in our integrated MPRA (*p*=0.0007), suggesting that integration of regulatory elements may allow for improved sensitivity. Next, we compared the activity of individual enhancers when tested in our integrated *Tnni1* MPRA, or an episomal MPRA with a *Tnni1* minimal promoter. Interestingly, the correlation between the two assays was poor (Fig. 4E), with similar poor correlations observed regardless of which minimal promoter was used in the episomal assay (Supp. Fig. 3E-H). Several enhancers exhibited high activity in only one of the assays, suggesting that variations in enhancer or locus architecture which are not yet understood may have a large impact on assay results. The correlation of P300 occupancy to enhancer activity was stronger in the episomal assay (Supp. Fig. 3I,J), further supporting this notion of mechanistic differences between assays.

## Discussion

CRISPR/Cas9-based genome editing technologies constitute a suite of powerful tools for gene manipulation. Many of these tools enable creation of targeted but imprecise edits (NHEJ), or very small precise edits (base editing). In this study, we demonstrate that CASAAV-HDR-based genome editing, which enables large, precise edits, can be employed *in vivo* within somatic cells for disease modeling and functional genomics. First, we used CASAAV-HDR to create and study two frameshifting titin variants that result in protein truncation and dilated cardiomyopathy in humans. By fusing a fluorescent tag to the C-termini of the truncated proteins, we were able to observe that the mutant proteins were stably expressed and at least partially incorporated into sarcomeres, although fluorescence unexpectedly localized near the Z-line.

In a second disease modeling application we utilized CASAAV-HDR to precisely delete arginine 14 of PLN - another mutation found in human cardiomyopathy patients. We found that PLN-R14Del did not dramatically disrupt protein localization or cell morphology, although a modest increase in cardiomyocyte width was observed. Interestingly, this pattern of concentric cardiomyocyte growth is typically observed in hypertrophic cardiomyopathy, while the R14Del mutation is linked to dilated cardiomyopathy. Our model system uniquely features assessment of the mutant cells in a genetic mosaic organ^60^, possibly revealing concentric cardiomyocyte hypertrophy as a cell autonomous phenotype while, due to organ failure, the manifested pattern in traditional models is eccentric cardiomyocyte hypertrophy. Our analysis of cardiomyocyte structure in six-week old mice is consistent with studies conducted in standard PLN-R14Del gene-targeted mice, which were reported after we had initiated our own work. In these studies, the effect of the mutation increases in severity over time, with PLN^R14Del/+^ cardiomyocytes being largely normal at eight weeks of age^38^. While we noted that cardiomyocyte architecture was largely unperturbed six weeks after editing, functional defects were more apparent. We observed marked inhibition of PLN phosphorylation, and an increase in peak amplitude variation that strongly resembled Ca^2+^ alternans. The PLN R14Del mutation results in SERCA inhibition, and SERCA inhibition is a known mechanism causing Ca^2+^ alternans^46^. Furthermore, action potential alternans have been previously associated with the R14Del mutation^61,62^. While our observation of Ca^2+^ alternans is therefore somewhat unsurprising, this appears to be the first report of Ca^2+^ alternans in a mammalian PLN R14Del disease model. More broadly, to our knowledge, this is the first study to use precise somatic genome editing to create and analyze tagged variants or to model disease variants in vivo within the mammalian heart.

In a third and final application we used CASAAV-HDR for functional genomics, demonstrating that enhancer pools can be precisely integrated at target loci, allowing for high-throughput measurement of their activities in postnatal cardiomyocytes. Here we used the native *Tnni1* gene, which has minimal expression in the adult stage, as the reporter gene in a STARR-seq experiment that featured enhancer insertion into the 3’ UTR. We assayed 50 pre-validated enhancers and observed a clear distinction between positive and negative controls, with high correlation between duplicate samples, suggesting that assaying much larger enhancer libraries is feasible. When compared to traditional AAV episome based MPRA results, our integrated assay displayed a significant but moderate improvement in dynamic range. While group averages only moderately differed, correlations between individual enhancers in integrated and episomal MPRAs were poor, suggesting that a thorough assessment of enhancer activity may require complementary episomal and integrated MPRAs. Our integrated assay demonstrates that the impact of *cis*-regulatory elements on expression of a native gene can be measured at high-throughput *in vivo* in a cell-type specific manner, and importantly, opens the door to additional novel functional genomics assays, such as those assessing the impact of *trans*-acting regulatory elements (ex. sgRNAs for pooled CRISPR screening) on the expression of a candidate gene.

CASAAV-HDR has some limitations. First, HDR efficiency varies broadly by target locus, with high expression loci exhibiting better editing efficiencies^13^. As a result, this technique is best used to edit strongly expressed genes, which may limit its utility. A second limitation is that CASAAV-HDR creates a range of outcomes in any edited tissue, including indel formation, edit heterozygosity, and edit homozygosity. Here we designed assays such that cells undergoing successful editing are labeled by mScarlet or an epitope tag, while unedited cells or cells with only indels are not tagged. However, we can not rule out the possibility that successfully tagged cells also have an allele with an indel. When possible we used edited control groups, and sgRNAs targeting non-coding sequences, such that indel formation is unlikely to result in a phenotype. When it is necessary to target coding sequence, such as when creating the Titin truncating variants, we limited our observations to only two areas, where a positive result is meaningful regardless of indel status: variant protein stability and localization to the sarcomere. A third limitation of CASAAV-HDR is that due to a need for high-dose donor template, the technique is only optimal for targeting one locus at a time, which precludes parallel evaluation of large numbers of precisely edited genes. However, the approach is compatible with assaying large numbers of gene variants at a single locus for disease modeling or protein engineering purposes, which is not possible with traditional germline gene targeting.

In summary, CASAAV-HDR is a rapid and powerful technical platform for precise gene editing in cardiomyocytes. The platform is well suited to a diversity of applications, as we have demonstrated for disease modeling and functional genomics. In the future we anticipate the development of many additional applications such as precise insertion of therapeutic transgenes at safe-harbor loci, in vivo clonal barcoding, and diverse gene perturbation screens.

## Acknowledgements

YZ was supported by NSFC82200265. WTP was supported by R01HL146634. NJV was supported by K99HL143194 and start-up funds from the Herman B Wells Center for Pediatric Research. Portions of this research were conducted on the Quartz High Performance Computing Cluster, which is supported in part by Lilly Endowment, Inc., through its support for the Indiana University Pervasive Technology Institute. Portions of this research were conducted on the O2 High Performance Computing Cluster, supported by the Research Computing Group, at Harvard Medical School. See http://rc.hms.harvard.edu for more information.

## Author Contributions

Y.Z., W.T.P. and N.J.V. conceived and designed the study. Y.Z. and N.J.V. executed experiments. J.S.K. generated plasmids, viruses, and other necessary reagents, and assisted with processing tissues and cells. J.M. and V.J.B. contributed to analysis of calcium handling data. Y.Z., Y.L., W.T.P. and N.J.V. analyzed and interpreted data. Y.Z, W.T.P. and N.J.V. wrote the paper.

## Data availability

The data in this publication have been deposited in the GEO database under accession code GSE280309. Both raw and processed data are available. P300 ChIP-sequencing data was mined from GSE195901 and GSE88789. Cardiomyocyte-specific gene expression across development was mined from GSE195901. A calcium analysis script, and scripts for analysis of MPRA datasets are available on GitHub at https://github.com/nvanduse/CASAAV-HDR_Applications

## Methods

### Mouse

All animal experiments were approved by the Institutional Animal Care and Use Committee of Boston Children’s Hospital. *R26^fsCas9-GFP/fsCas9-GFP^* mice^63^ and *Tnnt2-Cre* transgenic mice^64^ were acquired from Jackson Laboratories (Stock #026175 and #024240 respectively). *cTnnt2-Cre(+); R26^fscas9-GFP/+^* mice were generated by crossing *R26^fscas9-GFP/fscas9-GFP^* females with *cTnnt2-Cre(+)* male mice. The day new pups were detected was treated as postnatal day 0 (P0). PCR genotyping was conducted as previously reported^63,64^, via amplification of tail DNA isolated by overnight digestion with proteinase-k followed by precipitation with isopropanol.

### Plasmid

AAV-U6gRNA-5HA-2xHA-400SET-3HA, AAV-U6gRNA-5HA-mScarlet-3HA, and AAV-U6gRNA-5HA-Snap-3HA were constructed by restriction enzyme cloning or HiFi assembly. HDR vectors carried sequence for creation of mouse variants equivalent to human variants selected from the literature. In addition to PLN R14Del, mouse TTN-T65094fs was derived from the human variant p.Pro22582fs, and mouse TTN +AT63117fs was derived from the human variant 43628insAT.

Homology arms were PCR amplified from mouse genomic DNA, and inserted into each vector. Guide RNAs were synthesized as single-stranded oligos, annealed, and inserted into vectors cut by *BbsI*. mScarlet was synthesized as double-stranded DNA, and Snap sequence was PCR amplified from pAAV-myrSNAP (Addgene, #60098). Fragments were inserted into pAAV using T4 ligase (NEB, M0202L) or HiFi assembly kit (NEB, E5520S). For construction of AAV-U6gRNA-5HA-2xHA-3HA, 2xHA-3HA was PCR amplified from AAV-U6gRNA-5HA-P2A-mScarlet-3HA, then 2xHA-3HA was inserted into AAV-U6gRNA-5HA-P2A-mScarlet-3HA cut by *Not*l and *AsiS*l. Plasmid sequences can be found in Supp. Table 7.

### AAV9 vector production and injection

For each of ten 15 cm plates of HEK 293T cells, 7 μg of AAV2 genome (carrying transgene), 7 μg of AAV9 Rep/Cap, and 20 μg of pHGT1-adeno/dF6 helper were diluted with 1.8ml of Opti-Mem, and transfected with 170 μl of 1 μg/μl PEI (sigma 408727) transfection reagent. Cells were harvested 72 hours after transfection, and resuspended in AAV lysis buffer (20 mM Tris pH8.0, 1 mM MgCl_2_, 150 mM NaCl), and lysed by three freeze/thaw cycles. Media was also collected and precipitated overnight at 4°C by adding a 40% PEG 2M NaCl solution at a volume equal to 20% of the media volume. The precipitate was pelleted, resuspended in AAV lysis buffer, and combined with the cells. Cell genomic DNA was degraded by addition of 4ul of Pierce Universal Nuclease (Fisher PI88702) and incubation at 37’C for 20 minutes. AAV was purified by ultracentrifugation (45,000 RPM with rotor VTi 50, 110 minutes at 16℃) on an optiprep gradient (Cosmo Bio NC1059560). AAV in PBS + 0.001% F68 (Thermo Fisher, 24040032) was concentrated with an Amicon 100 kD column (UFC910024). AAV genomic DNA was isolated by proteinase K (Fisher BP1700100) digestion of purified vector, which was then titered by qPCR using an AAV-U6gRNA plasmid to make a standard curve. All mice were anesthetized with isoflurane prior to injection. Newborn mice were subcutaneously injected with 1×10˄12 vg of AAV-HDR vector.

### Isolation of cardiomyocytes

Isolation of cardiomyocytes was performed as previously described^65^. Briefly, mice were anesthetized with isoflurane and sacrificed by cervical dislocation. The heart was removed, and the aorta cannulated and fastened on a Langendorff apparatus with surgical thread. The heart was dissociated via retro-grade perfusion of the coronary arteries with collagenase type II, which was diluted in perfusion buffer (137 mM NaCl, 20 mM HEPES, 10 mM D-glucose, 5.4 mM KCl, 1.2 mM MgCl2, 1.2 mM NaH2PO4, 10 mM Taurine, 10 mM BDM) for 10 minutes. The heart was then transferred to a dish with stopping buffer (10% FBS + perfusion buffer), and gently dissociated into single cells via pipetting. Cells were passed through a 100 μm filter and cardiomyocytes were enriched by collecting the cell pellet after allowing cells to settle for 10 minutes by gravity.

### Immunostaining

Immunostaining was performed as previously described^65^. Briefly, the freshly isolated CMs were cultured in DMEM + 10 µM blebbistatin (EMD Millipore 203390) on laminin (Corning CB-40232) coated glass coverslips (VWR 100499-634), at 37℃ for 30 minutes to allow CMs to attach. CMs were then fixed with 4% paraformaldehyde in PBS (4% PFA/PBS) for 15 minutes at 37℃, rinsed with PBS for 2 times, followed with PBST (0.1% Triton-X 100 + PBS) for another 10 minutes at room temperature, washed with PBS for another 2 times, proceed with 4% BSA/PBS for blocking 10 minutes and incubated with the primary antibody (1:1000) in PBS for overnight at 4℃. The next day CMs were briefly rinsed with PBS for 3 times and incubated with the secondary antibody (1:500, Dapi is 1:100,000) in PBS for one hour at 4℃, followed by three PBS rinses. Coverslips with CMs were mounted on slides using Diamond Antifade mountant (Thermo, P36965) before imaging.

### Histology

Cardiac histology was performed as previously described^65^. Briefly, mice were euthanized with CO2 and the heart was removed and fixed with 4% PFA/PBS (4% paraformaldehyde + PBS) overnight at 4℃. Next, the fixed hearts were transferred to 15% sucrose/PBS for several hours, followed by 30% sucrose/PBS overnight at 4℃. The next day the hearts were embedded in tissue freezing medium (General Data, TFM-5) and frozen at-80℃. Heart tissue was cut into 10 micron sections using a cryostat (Thermo Scientific, Microm HM550). Samples were permeabilized and rehydrated in 0.1% PBST (0.1% Triton-X 100 + PBS) for 20 minutes, rinsed 2 times with PBS, blocked with 4% BSA/PBS for 1 hour, incubated with WGA (4 µg/ml) and Dapi for 1 hour, and rinsed with PBS twice. The sections coverslipped with Diamond Antifade mountant (Thermo, P36965) before imaging.

### Fluorescence imaging

We took fluorescent images by using confocal or Keyence. For in situ imaging, hearts were isolated from euthanized mice and cleaned by cold PBS. Then the heart was put on a glass-bottom dish, and immediately imaged by Olympus FV3000R inverted laser scanning confocal microscope with ×60 objective. For immunostaining CMs, fluorescent photos were taken by Olympus FV3000R inverted laser scanning confocal microscope with ×60 objective. For cryosections tissues, images were taken by Olympus FV3000R inverted laser scanning confocal microscope with ×30 or ×60 objective.

### T-tubule and Colocalization Analysis

Isolated cardiomyocytes immunostained with CAV3 were imaged on an Olympus FV3000R inverted laser scanning confocal microscope with 60x objective and 1.3x digital zoom, at 0.159 microns per pixel. Images were cropped to remove the cell border, leaving rectangular regions-of-interest (ROI). For T-tubule analysis, AutoTT was used to quantify CAV3 organization from the ROIs according to manual instructions^40^. For colocalization analysis, CAV3 and mScarlet-PLN signal within each ROI was analyzed using the ImageJ Coloc 2 plugin with default settings. Pearson correlations were calculated without thresholding.

### DNA and RNA extraction

For DNA isolation, the tail, heart tissue, or cardiomyocytes were digested with 0.5ml of DNA digestion buffer (50 mM Tris-HCl pH 8.0, 1 mM EDTA pH 8.0, 100 mM NaCl, 1% SDS) and 4 μl of 10 mg/ml proteinase K overnight at 55 ℃. The next day the genomic DNA was precipitated by isopropanol, rinsed with 70% ethanol, and suspended in water. For RNA isolation, 100ul of homogenized heart tissue or cardiomyocytes were added to 500ul of Trizol (Thermo Fisher Scientific, 15596018). Samples were vortexed, 150 ul of chloroform was added, vortexed again, and allowed to sit for 3 minutes at room temperature. Samples were then spun for 10 minutes at 12,000g, and the aqueous layer was transferred onto an RNA Clean & Concentrator Column and processed according to manufacturer instructions (Zymo research, R1016).

### Sanger sequencing of transcripts from edited loci

RNA was reverse transcribed into cDNA, and amplified in two rounds of PCR. In the first round a primer outside of the AAV homology arms was used to prevent amplification of any AAV genomic DNA or transcripts that may originate from the AAV. The second round of PCR featured primers nested inside the first set, resulting in production of a smaller amplicon that was submitted for Sanger sequencing.

### Recording calcium transients

Calcium was reintroduced into isolated CMs by transferring them through a series of perfusion buffer solutions with 60 μM, 240 μM, 600 μM, and 1.2 mM calcium. For each step, cells were gently resuspended and allowed to settle naturally for ∼10 minutes, at which point the supernatant was removed and the next solution was added. After the last calcium reintroduction step, cells were cultured in a 96 well-plate with MEM media containing 2 μM Rhod-2, 0.6 μM SNAP-Cell^®^ 647-SiR, and Hoechst (1:1000) at 37 ℃ for 30 minutes. Next, cells were washed twice with Tyrode’s solution (140mM Nacl, 4mM KCl, 1.2mM CaCl_2_, 2mM MgCl_2_, 10 mM HEPES, 15 mM Glucose, 2 mM Pyruvate, pH=7.4). Then, the cells were transferred into the 37 ℃ chamber of the Kinetic Image Cytometer (Vala Science) and electrically stimulated (2Hz at 30 voltage; paced for 30 cycles with recording of the last 5). SNAP-tag signal and temporal calcium signals were acquired for individual cells and quantified with ImageJ.

### Analysis of calcium transients

Standard transient characteristics were calculated as previously described^66^, from five cycles. Briefly, cells were manually segmented, and the mean intensity of each segmented cell over time was input into a custom MATLAB (Mathworks) script for data filtering and calcium transient parameter calculation. Data filtration was performed by removing photobleaching and applying a median filter.

After normalizing data to baseline and identifying peaks via the MATLAB findpeaks function, parameter calculations included amplitude, maximal upstroke velocity, maximal downstroke velocity, and time from peak to 50% decay in calcium transient. For analysis of calcium abnormalities, unaveraged traces of the five recorded cycles were uploaded to AnomalyExplorer. For a small number of cells, technical problems resulted in premature termination of recording prior to acquisition of five full cycles; these cells were excluded. AnomalyExplorer allows for data exploration via real-time visualization of anomalies across adjustable analysis specifications. We explored the frequency of the various anomaly types detected by AnomalyExplorer across a range of specifications, and noted an increase of “low peaks”. This was most apparent when normal peak height was stringently defined (low peaks = 10-94%; Fig. 3H). Visual inspection of the data suggested there was an overall increase in amplitude variation, rather than a small number of low peaks, so we directly measured variation by extracting calcium peak coordinates for each cardiomyocyte, fitting them with a best fit line, and measuring the distance of each peak from the line. This distance was normalized to peak height and plotted. Code used in the analysis is available at https://github.com/nvanduse/CASAAV-HDR_Applications.

### Western Blot

P42 mice were intraperitoneally injected with isoproterenol (5 μg/g) for 5 minutes before euthanasia, hearts were homogenized in 500 μl protein lysis buffer (120mM NaCl, 40mM HEPES, 1 mM EDTA, 10 mM Glycerophosphate, 0.3% CHAPS, 1% Triton X-100, 1× protease/phosphatase inhibitor cocktail) and sonicated for 90s. Lysates were spun at 12,000g for 15 minutes at 4 ℃, and the supernatant transferred to a new tube. Protein concentration was measured by BCA Protein Assay Kit. 0.2 mg protein of each sample together with primary antibody (PLN and pPLN are 1: 250, GAPDH is 1:400) and anti-rabbit secondary antibody were added into an 8 × 25 capillary cartridge, inserted into the Wes machine, and used to detect target proteins.

### Genomically integrated MPRA

A total of fifty-five enhancers with activity in cardiomyocytes (25), cardiac endothelial cells (25), or mouse embryonic stem cells (5), were selected from the literature or the VISTA Enhancer Database (Supp. Table 4). P300 ChIP-seq datasets from each cell type were mined and analyzed for each full length enhancer, and a 400 bp interval most overlapping with P300 occupancy was selected. These 400 bp regions, along with a 3’ barcode, were synthesized by Twist Biosciences. Synthesized enhancers were ligated into the Tnni1 HDR donor vector (Supp. Table 6), and sequenced to confirm successful cloning. One endothelial enhancer was missing, possibly due to failed synthesis. We proceeded to package the plasmid pool into AAV9, and injected 1×10˄12 vg into newborn cTnnt2-Cre(+); R26^fsCas9-P2A-GFP^ mice, as described above. Hearts were collected at P28 and dissociated into single cell suspensions. A small fraction of cardiomyocytes from each heart was seeded in DMEM, fixed, and stained for the HA-tag. The remaining cardiomyocytes for each heart were divided into two portions, with genomic DNA being isolated from one portion and RNA isolated from the other, as described above. For preparation of amplicon-seq libraries from RNA, for each mouse, duplicate 2ug RNA samples were reverse transcribed using Oligo-dT with SMARTScribe Reverse Transcriptase (Takara 639537) following the manufacturer instructions. For each cDNA sample, 5ul of cDNA was amplified in duplicate Phusion Hot Start Flex (NEB M0536L) reactions, with a touchdown protocol for 15 cycles, decreasing 1°C per cycle from 70°C to 55°C annealing temperature. The forward primer targeted a priming site situated between the enhancer and barcode, while the reverse primer targeted the Tnni3 UTR downstream of the HDR-vector homology arm (Fig. 4B, red arrows). The forward primer added a full length NGS adapter via primer tailing, while the reverse primer added a partial adapter. In a second round of PCR, 2ul of first round product was amplified in duplicate for 20 cycles with the same forward primer and an indexed reverse primer that completed addition of the reverse NGS adapter. After combining replicates, 5ul of each sample was run on a gel, and the amplicon concentration was estimated via densitometry. Samples were then combined in equal proportions, run together on a gel, and the amplicon was extracted, purified, and submitted for amplicon-sequencing. The same approach, excluding reverse transcription which is not applicable, was used to prepare amplicons from mouse genomic DNA, with the following differences: 1) The forward primer for the first round of amplification targeted an upstream region of the Tnni1 gene, outside the 5’ homology arm, while the reverse primer targeted a region just downstream of the insertion, within the 3’ homology arm, and added a partial NGS adapter (Fig. 4B, black arrows). 2) In the second round of amplification the forward primer targeted the insertion priming site located between the enhancer and barcode (same primer as for RNA), while the reverse primer added a multiplexing index and completed the adapter. See Supplemental table 1 for primer sequences. NGS sequences were trimmed to remove non-barcode sequences, and then the number of times each barcode occurred in each sample was quantified. These readcounts were used to calculate reads per million (RPM), and activity was calculated as the ratio between barcode frequency in RNA versus DNA. Normalized activity (for comparison to episomal MPRA data) was calculated by multiplying activity data by an adjustment factor, such that the average of the endothelial enhancer group was equal to one. NGS processing script is available on Github (see data availability statement).

### Episomal MPRAs

Barcoded enhancers and promoters were cloned into the MPRA vector as shown in Fig. S3D. The plasmid pool was used to produce a single AAV9 library, which was subcutaneously injected into six newborn mice with cardiomyocyte specific activation of Cas9, as described above. Hearts were collected at P28 and RNA was isolated from whole ventricles. RNA was reverse transcribed with Oligo-dT priming. For DNA samples, DNA was isolated from untransduced AAV via Proteinase-K digestion of the viral capsid, in three technical replicates, as previously described^21,65^. RNA and DNA samples were amplified in two rounds of PCR (see Supplemental table 1 for primers), and submitted for amplicon-sequencing as described above. Promoter--enhancer barcode combinations were quantified, and normalized for sequencing depth (CPM). Normalized activity was calculated as described above for all enhancer-promoter pairs that were present at >30 CPM in all DNA replicates. NGS processing script is available on Github (see data availability statement).

## References

1. Barrangou R, Fremaux C, Deveau H, Richards M, Boyaval P, Moineau S, Romero DA, Horvath P. CRISPR provides acquired resistance against viruses in prokaryotes. Science. 2007 Mar 23;315(5819):1709–1712. PMID: 17379808

2. Cong L, Ran FA, Cox D, Lin S, Barretto R, Habib N, Hsu PD, Wu X, Jiang W, Marraffini LA, Zhang F. Multiplex genome engineering using CRISPR/Cas systems. Science. 2013 Feb 15;339(6121):819–823. PMCID: PMC3795411

3. Jinek M, Chylinski K, Fonfara I, Hauer M, Doudna JA, Charpentier E. A programmable dual-RNA-guided DNA endonuclease in adaptive bacterial immunity. Science. 2012 Aug 17;337(6096):816–821. PMCID: PMC6286148

4. Wang H, Yang H, Shivalila CS, Dawlaty MM, Cheng AW, Zhang F, Jaenisch R. One-step generation of mice carrying mutations in multiple genes by CRISPR/Cas-mediated genome engineering. Cell. 2013 May 9;153(4):910–918. PMCID: PMC3969854

5. Mali P, Yang L, Esvelt KM, Aach J, Guell M, DiCarlo JE, Norville JE, Church GM. RNA-guided human genome engineering via Cas9. Science. 2013 Feb 15;339(6121):823–826. PMCID: PMC3712628

6. Sander JD, Joung JK. CRISPR-Cas systems for editing, regulating and targeting genomes. Nat Biotechnol. 2014 Apr;32(4):347–355. PMCID: PMC4022601

7. Rothkamm K, Krüger I, Thompson LH, Löbrich M. Pathways of DNA double-strand break repair during the mammalian cell cycle. Mol Cell Biol. 2003 Aug;23(16):5706–5715. PMCID: PMC166351

8. VanDusen NJ, Guo Y, Gu W. CASAAV: A CRISPR-Based Platform for Rapid Dissection of Gene Function In Vivo. Current protocols in [Internet]. Wiley Online Library; 2017; Available from: https://currentprotocols.onlinelibrary.wiley.com/doi/abs/10.1002/cpmb.46

9. Guo Y, VanDusen NJ, Zhang L, Gu W, Sethi I, Guatimosim S, Ma Q, Jardin BD, Ai Y, Zhang D, Chen B, Guo A, Yuan GC, Song LS, Pu WT. Analysis of Cardiac Myocyte Maturation Using CASAAV, a Platform for Rapid Dissection of Cardiac Myocyte Gene Function In VivoNovelty and Significance. Circulation. Am Heart Assoc; 2017 Jun 9;120(12):1874–1888. PMCID: PMC5466492

10. Sarin S, Zuniga-Sanchez E, Kurmangaliyev YZ, Cousins H, Patel M, Hernandez J, Zhang KX, Samuel MA, Morey M, Sanes JR, Zipursky SL. Role for Wnt Signaling in Retinal Neuropil Development: Analysis via RNA-Seq and In Vivo Somatic CRISPR Mutagenesis. Neuron. 2018 Apr 4;98(1):109–126.e8. PMCID: PMC5930001

11. Martin A, Serano JM, Jarvis E, Bruce HS, Wang J, Ray S, Barker CA, O’Connell LC, Patel NH. CRISPR/Cas9 Mutagenesis Reveals Versatile Roles of Hox Genes in Crustacean Limb Specification and Evolution. Curr Biol. 2016 Jan 11;26(1):14–26. PMID: 26687626

12. Cox DBT, Platt RJ, Zhang F. Therapeutic genome editing: prospects and challenges. Nat Med. 2015 Feb;21(2):121–131. PMCID: PMC4492683

13. Zheng Y, VanDusen NJ, Butler CE, Ma Q, King JS, Pu WT. Efficient In Vivo Homology-Directed Repair Within Cardiomyocytes. Circulation. Am Heart Assoc; 2022 Mar 8;145(10):787–789. PMID: 35254919

14. Hinson JT, Chopra A, Nafissi N, Polacheck WJ, Benson CC, Swist S, Gorham J, Yang L, Schafer S, Sheng CC, Haghighi A, Homsy J, Hubner N, Church G, Cook SA, Linke WA, Chen CS, Seidman JG, Seidman CE. HEART DISEASE. Titin mutations in iPS cells define sarcomere insufficiency as a cause of dilated cardiomyopathy. Science. 2015 Aug 28;349(6251):982–986. PMCID: PMC4618316

15. Gerull B, Gramlich M, Atherton J, McNabb M, Trombitás K, Sasse-Klaassen S, Seidman JG, Seidman C, Granzier H, Labeit S, Frenneaux M, Thierfelder L. Mutations of TTN, encoding the giant muscle filament titin, cause familial dilated cardiomyopathy. Nat Genet. 2002 Feb;30(2):201–204. PMID: 11788824

16. Haghighi K, Kolokathis F, Gramolini AO, Waggoner JR, Pater L, Lynch RA, Fan GC, Tsiapras D, Parekh RR, Dorn GW 2nd, MacLennan DH, Kremastinos DT, Kranias EG. A mutation in the human phospholamban gene, deleting arginine 14, results in lethal, hereditary cardiomyopathy. Proc Natl Acad Sci U S A. 2006 Jan 31;103(5):1388–1393. PMCID: PMC1360586

17. Karakikes I, Stillitano F, Nonnenmacher M, Tzimas C, Sanoudou D, Termglinchan V, Kong CW, Rushing S, Hansen J, Ceholski D, Kolokathis F, Kremastinos D, Katoulis A, Ren L, Cohen N, Gho JMIH, Tsiapras D, Vink A, Wu JC, Asselbergs FW, Li RA, Hulot JS, Kranias EG, Hajjar RJ. Correction of human phospholamban R14del mutation associated with cardiomyopathy using targeted nucleases and combination therapy. Nat Commun. 2015 Apr 29;6:6955. PMCID: PMC4421839

18. Haghighi K, Pritchard T, Bossuyt J, Waggoner JR, Yuan Q, Fan GC, Osinska H, Anjak A, Rubinstein J, Robbins J, Bers DM, Kranias EG. The human phospholamban Arg14-deletion mutant localizes to plasma membrane and interacts with the Na/K-ATPase. J Mol Cell Cardiol. 2012 Mar;52(3):773–782. PMCID: PMC3376549

19. Jiang X, Xu Y, Sun J, Wang L, Guo X, Chen Y. The phenotypic characteristic observed by cardiac magnetic resonance in a PLN-R14del family. Sci Rep. 2020 Oct 5;10(1):16478. PMCID: PMC7536202

20. Akerberg BN, Gu F, VanDusen NJ, Zhang X, Dong R, Li K, Zhang B, Zhou B, Sethi I, Ma Q, Wasson L, Wen T, Liu J, Dong K, Conlon FL, Zhou J, Yuan GC, Zhou P, Pu WT. A reference map of murine cardiac transcription factor chromatin occupancy identifies dynamic and conserved enhancers. Nat Commun. nature.com; 2019 Oct 28;10(1):4907. PMCID: PMC6817842

21. Zhou P, VanDusen NJ, Zhang Y, Cao Y, Sethi I, Hu R, Zhang S, Wang G, Ye L, Mazumdar N, Chen J, Zhang X, Guo Y, Li B, Ma Q, Lee JY, Gu W, Yuan GC, Ren B, Chen K, Pu WT. Dynamic changes in P300 enhancers and enhancer-promoter contacts control mouse cardiomyocyte maturation. Dev Cell. 2023 May 22;58(10):898–914.e7. PMCID: PMC10231645

22. Cao Y, Zhang X, Akerberg BN, Yuan H, Sakamoto T, Xiao F, VanDusen NJ, Zhou P, Sweat ME, Wang Y, Prondzynski M, Chen J, Zhang Y, Wang P, Kelly DP, Pu WT. In Vivo Dissection of Chamber-Selective Enhancers Reveals Estrogen-Related Receptor as a Regulator of Ventricular Cardiomyocyte Identity. Circulation [Internet]. Am Heart Assoc; 2023 Jan 27; Available from: 10.1161/CIRCULATIONAHA.122.061955 PMID: 36705030

23. Muerdter F, Boryń ŁM, Arnold CD. STARR-seq - principles and applications. Genomics. 2015 Sep;106(3):145–150. PMID: 26072434

24. Gaynor R. Cellular transcription factors involved in the regulation of HIV-1 gene expression. AIDS. 1992 Apr;6(4):347–363. PMID: 1616633

25. Pereira LA, Bentley K, Peeters A, Churchill MJ, Deacon NJ. SURVEY AND SUMMARY A compilation of cellular transcription factor interactions with the HIV-1 LTR promoter. Nucleic Acids Res. Oxford Academic; 2000 Feb 1;28(3):663–668.

26. Julien L, Chassagne J, Peccate C, Lorain S, Piétri-Rouxel F, Danos O, Benkhelifa-Ziyyat S. RFX1 and RFX3 Transcription Factors Interact with the D Sequence of Adeno-Associated Virus Inverted Terminal Repeat and Regulate AAV Transduction. Sci Rep. 2018 Jan 9;8(1):210. PMCID: PMC5760533

27. Earley LF, Conatser LM, Lue VM, Dobbins AL, Li C, Hirsch ML, Samulski RJ. Adeno-Associated Virus Serotype-Specific Inverted Terminal Repeat Sequence Role in Vector Transgene Expression. Hum Gene Ther. 2020 Feb;31(3-4):151–162. PMCID: PMC7047122

28. Delenda C. Lentiviral vectors: optimization of packaging, transduction and gene expression. J Gene Med. 2004 Feb;6 Suppl 1:S125–38. PMID: 14978756

29. Herman DS, Lam L, Taylor MRG, Wang L, Teekakirikul P, Christodoulou D, Conner L, DePalma SR, McDonough B, Sparks E, Teodorescu DL, Cirino AL, Banner NR, Pennell DJ, Graw S, Merlo M, Di Lenarda A, Sinagra G, Bos JM, Ackerman MJ, Mitchell RN, Murry CE, Lakdawala NK, Ho CY, Barton PJR, Cook SA, Mestroni L, Seidman JG, Seidman CE. Truncations of titin causing dilated cardiomyopathy. N Engl J Med. 2012 Feb 16;366(7):619–628. PMCID: PMC3660031

30. Yotti R, Seidman CE, Seidman JG. Advances in the Genetic Basis and Pathogenesis of Sarcomere Cardiomyopathies. Annu Rev Genomics Hum Genet. 2019 Aug 31;20:129–153. PMID: 30978303

31. Roberts AM, Ware JS, Herman DS, Schafer S, Baksi J, Bick AG, Buchan RJ, Walsh R, John S, Wilkinson S, Mazzarotto F, Felkin LE, Gong S, MacArthur JAL, Cunningham F, Flannick J, Gabriel SB, Altshuler DM, Macdonald PS, Heinig M, Keogh AM, Hayward CS, Banner NR, Pennell DJ, O’Regan DP, San TR, de Marvao A, Dawes TJW, Gulati A, Birks EJ, Yacoub MH, Radke M, Gotthardt M, Wilson JG, O’Donnell CJ, Prasad SK, Barton PJR, Fatkin D, Hubner N, Seidman JG, Seidman CE, Cook SA. Integrated allelic, transcriptional, and phenomic dissection of the cardiac effects of titin truncations in health and disease. Sci Transl Med. 2015 Jan 14;7(270):270ra6. PMCID: PMC4560092

32. Tharp CA, Haywood ME, Sbaizero O, Taylor MRG, Mestroni L. The Giant Protein Titin’s Role in Cardiomyopathy: Genetic, Transcriptional, and Post-translational Modifications of TTN and Their Contribution to Cardiac Disease. Front Physiol. 2019 Nov 28;10:1436. PMCID: PMC6892752

33. Schafer S, de Marvao A, Adami E, Fiedler LR, Ng B, Khin E, Rackham OJL, van Heesch S, Pua CJ, Kui M, Walsh R, Tayal U, Prasad SK, Dawes TJW, Ko NSJ, Sim D, Chan LLH, Chin CWL, Mazzarotto F, Barton PJ, Kreuchwig F, de Kleijn DPV, Totman T, Biffi C, Tee N, Rueckert D, Schneider V, Faber A, Regitz-Zagrosek V, Seidman JG, Seidman CE, Linke WA, Kovalik JP, O’Regan D, Ware JS, Hubner N, Cook SA. Titin-truncating variants affect heart function in disease cohorts and the general population. Nat Genet. 2017 Jan;49(1):46–53. PMCID: PMC5201198

34. Tskhovrebova L, Trinick J. Roles of titin in the structure and elasticity of the sarcomere. J Biomed Biotechnol. 2010 Jun 21;2010:612482. PMCID: PMC2896707

35. da Silva Lopes K, Pietas A, Radke MH, Gotthardt M. Titin visualization in real time reveals an unexpected level of mobility within and between sarcomeres. J Cell Biol. 2011 May 16;193(4):785–798. PMCID: PMC3166869

36. MacLennan DH, Kranias EG. Phospholamban: a crucial regulator of cardiac contractility. Nat Rev Mol Cell Biol. 2003 Jul;4(7):566–577. PMID: 12838339

37. Vafiadaki E, Haghighi K, Arvanitis DA, Kranias EG, Sanoudou D. Aberrant PLN-R14del Protein Interactions Intensify SERCA2a Inhibition, Driving Impaired Ca2+ Handling and Arrhythmogenesis. Int J Mol Sci [Internet]. 2022 Jun 22;23(13). Available from: 10.3390/ijms23136947 PMCID: PMC9266971

38. Eijgenraam TR, Boukens BJ, Boogerd CJ, Schouten EM, van de Kolk CWA, Stege NM, Te Rijdt WP, Hoorntje ET, van der Zwaag PA, van Rooij E, van Tintelen JP, van den Berg MP, van der Meer P, van der Velden J, Silljé HHW, de Boer RA. The phospholamban p.(Arg14del) pathogenic variant leads to cardiomyopathy with heart failure and is unreponsive to standard heart failure therapy. Sci Rep. 2020 Jun 17;10(1):9819. PMCID: PMC7300032

39. Guo A, Zhang C, Wei S, Chen B, Song LS. Emerging mechanisms of T-tubule remodelling in heart failure. Cardiovasc Res. 2013 May 1;98(2):204–215. PMCID: PMC3697065

40. Guo A, Song LS. AutoTT: automated detection and analysis of T-tubule architecture in cardiomyocytes. Biophys J. Elsevier; 2014 Jun 17;106(12):2729–2736. PMCID: PMC4070273

41. Brittsan AG, Carr AN, Schmidt AG, Kranias EG. Maximal inhibition of SERCA2 Ca(2+) affinity by phospholamban in transgenic hearts overexpressing a non-phosphorylatable form of phospholamban. J Biol Chem. 2000 Apr 21;275(16):12129–12135. PMID: 10766848

42. Ceholski DK, Trieber CA, Holmes CFB, Young HS. Lethal, hereditary mutants of phospholamban elude phosphorylation by protein kinase A. J Biol Chem. 2012 Aug 3;287(32):26596–26605. PMCID: PMC3411000

43. Landstrom AP, Dobrev D, Wehrens XHT. Calcium Signaling and Cardiac Arrhythmias. Circ Res. 2017 Jun 9;120(12):1969–1993. PMCID: PMC5607780

44. Penttinen K, Siirtola H, Àvalos-Salguero J, Vainio T, Juhola M, Aalto-Setälä K. Novel Analysis Software for Detecting and Classifying Ca2+ Transient Abnormalities in Stem Cell-Derived Cardiomyocytes. PLoS One. 2015 Aug 26;10(8):e0135806. PMCID: PMC4550257

45. Short B. Understanding Ca2+ alternans. J Gen Physiol [Internet]. 2021 Feb 1;153(2). Available from: 10.1085/jgp.202112862 PMCID: PMC7812828

46. Millet J, Aguilar-Sanchez Y, Kornyeyev D, Bazmi M, Fainstein D, Copello JA, Escobar AL. Thermal modulation of epicardial Ca2+ dynamics uncovers molecular mechanisms of Ca2+ alternans. J Gen Physiol [Internet]. 2021 Feb 1;153(2). Available from: 10.1085/jgp.202012568 PMCID: PMC7797898

47. Qu Z, Liu MB, Nivala M. A unified theory of calcium alternans in ventricular myocytes. Sci Rep. 2016 Oct 20;6:35625. PMCID: PMC5071909

48. Visel A, Minovitsky S, Dubchak I, Pennacchio LA. VISTA Enhancer Browser--a database of tissue-specific human enhancers. Nucleic Acids Res. 2007 Jan;35(Database issue):D88–92. PMCID: PMC1716724

49. Zhou P, Gu F, Zhang L, Akerberg BN, Ma Q, Li K, He A, Lin Z, Stevens SM, Zhou B, Pu WT. Mapping cell type-specific transcriptional enhancers using high affinity, lineage-specific Ep300 bioChIP-seq. Elife [Internet]. elifesciences.org; 2017 Jan 25;6. Available from: 10.7554/eLife.22039 PMCID: PMC5295818

50. Okada Y, Yano K, Jin E, Funahashi N, Kitayama M, Doi T, Spokes K, Beeler DL, Shih SC, Okada H, Danilov TA, Maynard E, Minami T, Oettgen P, Aird WC. A three-kilobase fragment of the human Robo4 promoter directs cell type-specific expression in endothelium. Circ Res. 2007 Jun 22;100(12):1712–1722. PMID: 17495228

51. Schlaeger TM, Bartunkova S, Lawitts JA, Teichmann G, Risau W, Deutsch U, Sato TN. Uniform vascular-endothelial-cell-specific gene expression in both embryonic and adult transgenic mice. Proc Natl Acad Sci U S A. 1997 Apr 1;94(7):3058–3063. PMCID: PMC20321

52. De Val S, Chi NC, Meadows SM, Minovitsky S, Anderson JP, Harris IS, Ehlers ML, Agarwal P, Visel A, Xu SM, Pennacchio LA, Dubchak I, Krieg PA, Stainier DYR, Black BL. Combinatorial regulation of endothelial gene expression by ets and forkhead transcription factors. Cell. 2008 Dec 12;135(6):1053–1064. PMCID: PMC2782666

53. Göttgens B, Broccardo C, Sanchez MJ, Deveaux S, Murphy G, Göthert JR, Kotsopoulou E, Kinston S, Delaney L, Piltz S, Barton LM, Knezevic K, Erber WN, Begley CG, Frampton J, Green AR. The scl +18/19 stem cell enhancer is not required for hematopoiesis: identification of a 5’ bifunctional hematopoietic-endothelial enhancer bound by Fli-1 and Elf-1. Mol Cell Biol. 2004 Mar;24(5):1870–1883. PMCID: PMC350551

54. Wythe JD, Dang LTH, Devine WP, Boudreau E, Artap ST, He D, Schachterle W, Stainier DYR, Oettgen P, Black BL, Bruneau BG, Fish JE. ETS factors regulate Vegf-dependent arterial specification. Dev Cell. 2013 Jul 15;26(1):45–58. PMCID: PMC3754838

55. Khandekar M, Brandt W, Zhou Y, Dagenais S, Glover TW, Suzuki N, Shimizu R, Yamamoto M, Lim KC, Engel JD. A Gata2 intronic enhancer confers its pan-endothelia-specific regulation. Development. 2007 May;134(9):1703–1712. PMID: 17395646

56. De Val S, Anderson JP, Heidt AB, Khiem D, Xu SM, Black BL. Mef2c is activated directly by Ets transcription factors through an evolutionarily conserved endothelial cell-specific enhancer. Dev Biol. 2004 Nov 15;275(2):424–434. PMID: 15501228

57. Wu J, Iwata F, Grass JA, Osborne CS, Elnitski L, Fraser P, Ohneda O, Yamamoto M, Bresnick EH. Molecular determinants of NOTCH4 transcription in vascular endothelium. Mol Cell Biol. 2005 Feb;25(4):1458–1474. PMCID: PMC548019

58. Donaldson IJ, Chapman M, Kinston S, Landry JR, Knezevic K, Piltz S, Buckley N, Green AR, Göttgens B. Genome-wide identification of cis-regulatory sequences controlling blood and endothelial development. Hum Mol Genet. 2005 Mar 1;14(5):595–601. PMID: 15649946

59. Becker PW, Sacilotto N, Nornes S, Neal A, Thomas MO, Liu K, Preece C, Ratnayaka I, Davies B, Bou-Gharios G, De Val S. An Intronic Flk1 Enhancer Directs Arterial-Specific Expression via RBPJ-Mediated Venous Repression. Arterioscler Thromb Vasc Biol. 2016 Jun;36(6):1209–1219. PMCID: PMC4894770

60. Guo Y, Pu WT. Genetic Mosaics for Greater Precision in Cardiovascular Research. Circ Res. 2018 Jun 22;123(1):27–29. PMCID: PMC6025841

61. Kamel SM, van Opbergen CJM, Koopman CD, Verkerk AO, Boukens BJD, de Jonge B, Onderwater YL, van Alebeek E, Chocron S, Polidoro Pontalti C, Weuring WJ, Vos MA, de Boer TP, van Veen TAB, Bakkers J. Istaroxime treatment ameliorates calcium dysregulation in a zebrafish model of phospholamban R14del cardiomyopathy. Nat Commun. 2021 Dec 9;12(1):7151. PMCID: PMC8660846

62. Raad N, Bittihn P, Cacheux M, Jeong D, Ilkan Z, Ceholski D, Kohlbrenner E, Zhang L, Cai CL, Kranias EG, Hajjar RJ, Stillitano F, Akar FG. Arrhythmia Mechanism and Dynamics in a Humanized Mouse Model of Inherited Cardiomyopathy Caused by Phospholamban R14del Mutation. Circulation. 2021 Aug 10;144(6):441–454. PMCID: PMC8456417

63. Platt RJ, Chen S, Zhou Y, Yim MJ, Swiech L, Kempton HR, Dahlman JE, Parnas O, Eisenhaure TM, Jovanovic M, Graham DB, Jhunjhunwala S, Heidenreich M, Xavier RJ, Langer R, Anderson DG, Hacohen N, Regev A, Feng G, Sharp PA, Zhang F. CRISPR-Cas9 knockin mice for genome editing and cancer modeling. Cell. 2014 Oct 9;159(2):440–455. PMCID: PMC4265475

64. Jiao K, Kulessa H, Tompkins K, Zhou Y, Batts L, Baldwin HS, Hogan BLM. An essential role of Bmp4 in the atrioventricular septation of the mouse heart. Genes Dev. 2003 Oct 1;17(19):2362–2367. PMCID: PMC218073

65. VanDusen NJ, Lee JY, Gu W, Butler CE, Sethi I, Zheng Y, King JS, Zhou P, Suo S, Guo Y, Ma Q, Yuan GC, Pu WW. Massively parallel in vivo CRISPR screening identifies RNF20/40 as epigenetic regulators of cardiomyocyte maturation. Nat Commun. 2021 Jul 21;12(1):4442. PMID: 34290256

66. Reyes Gaido OE, Pavlaki N, Granger JM, Mesubi OO, Liu B, Lin BL, Long A, Walker D, Mayourian J, Schole KL, Terrillion CE, Nkashama LJ, Hulsurkar MM, Dorn LE, Ferrero KM, Huganir RL, Müller FU, Wehrens XHT, Liu JO, Luczak ED, Bezzerides VJ, Anderson ME. An improved reporter identifies ruxolitinib as a potent and cardioprotective CaMKII inhibitor. Sci Transl Med. 2023 Jun 21;15(701):eabq7839. PMCID: PMC11022683

