## Supplemental Figures and Tables for "Cardiac Applications of CRISPR/AAV-Mediated Precise Genome Editing"

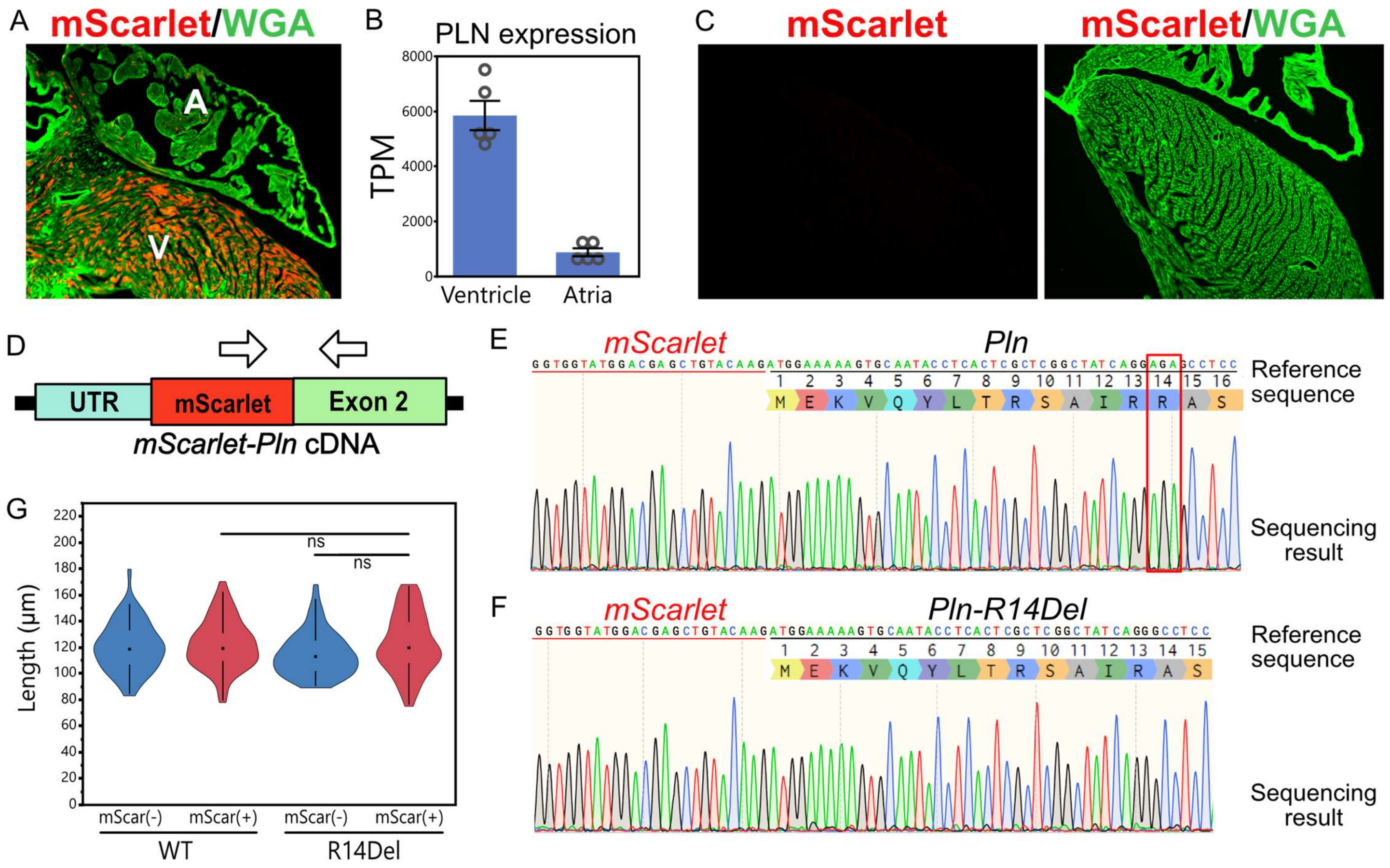


**Suppl. Fig. 1. Generation and analysis of PLN R14Del cardiomyocytes A.** Tissue section showing enriched mScarlet fluorescence in the ventricles (V) of PLN-WT hearts relative to the atria (A). **B.** Chamber specific *Pln* transcription in wildtype neonatal cardiomyocytes; mined from Cao et al. 2023. **C.** Cardiac tissue section from a wildtype (Cas9 negative) mouse injected with PLN-WT vector. **D.** Schematic of mScarlet/PLN junction to be Sanger sequenced from edited cDNA. **E.** Junction Sanger sequencing for PLN-WT hearts. Red box highlights R14. **F.** Junction Sanger sequencing for PLN-R14Del hearts. **G.** Length of isolated adult cardiomyocytes. Dunnett's *p* multiple comparison to R14Del mScarlet(+) group. ns, not significant. n = 56, 63, 63, and 62, for WT SN-, WT SN+, R14Del SN-, and R14Del SN+, respectively.

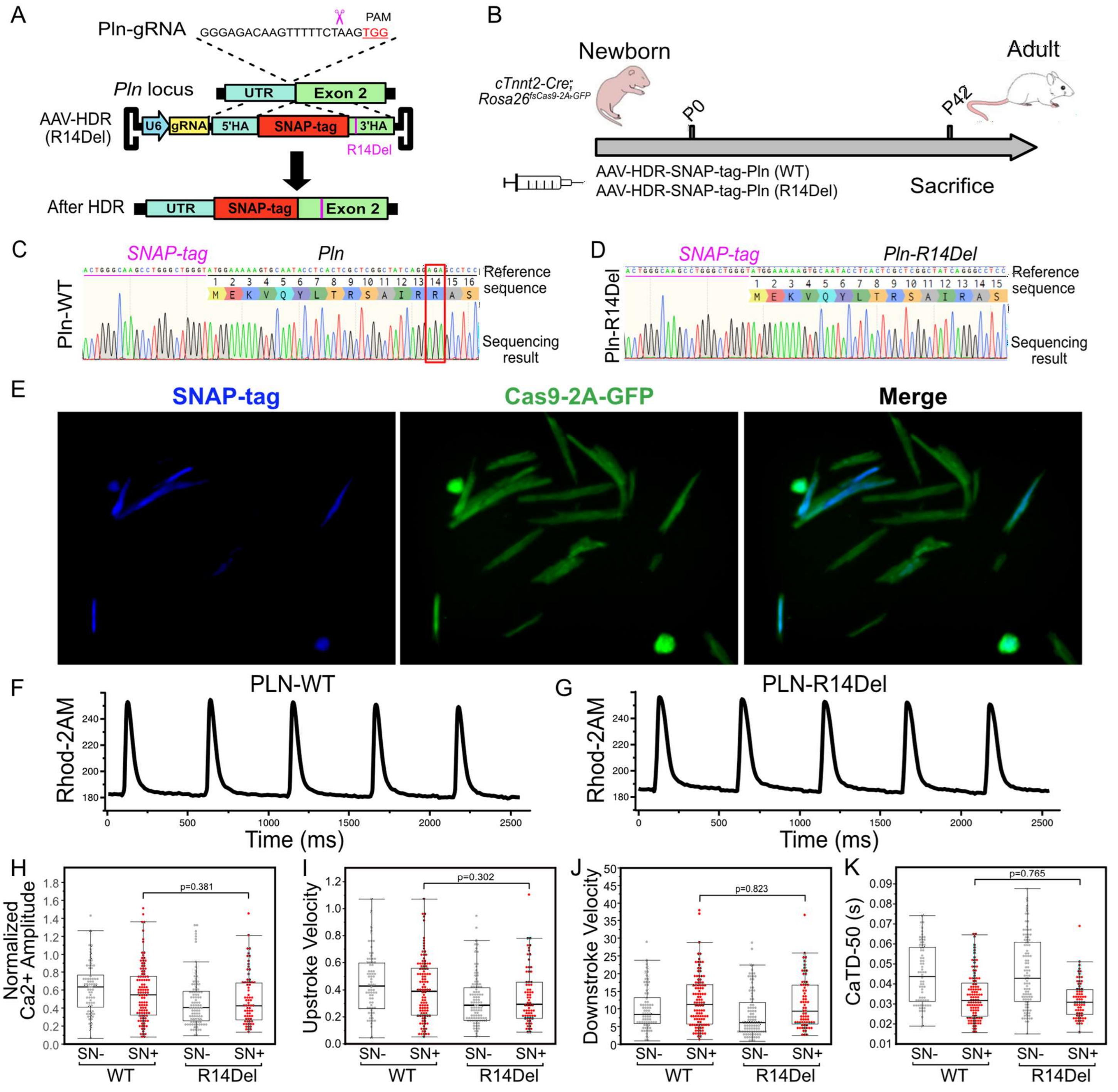


**Suppl. Fig. 2. Calcium transient analysis in PLN R14Del cardiomyocytes. A.** Homology directed repair-based strategy for fusing a SNAP-tag to wildtype and R14Del PLN. **B.** Experimental timeline. **C.** Sanger sequencing of cDNA from SNAP-Pln edited allele. **D.** Sanger sequencing of cDNA from SNAP-Pln-R14Del edited allele. **E.** SNAP-tag based far-red labeling of edited cardiomyocytes. Rhod-2AM labeled calcium transients were imaged in the red channel (not shown). **F.** Representative calcium trace for a PLN-WT cardiomyocyte. **G.** Representative calcium trace for a PLN-R14Del cardiomyocyte. **H.** Calcium transient peak amplitude. **I.** Calcium transient upstroke velocity. **J.** Calcium transient downstroke velocity. **K.** Action potential duration at 50% of the peak. Dunnett's *p* multiple comparison to R14Del mScarlet(+) group. For panels H-K, WT SN-, WT SN+, R14Del SN-, and R14Del SN+ n = 70, 106, 98, and 70, respectively.

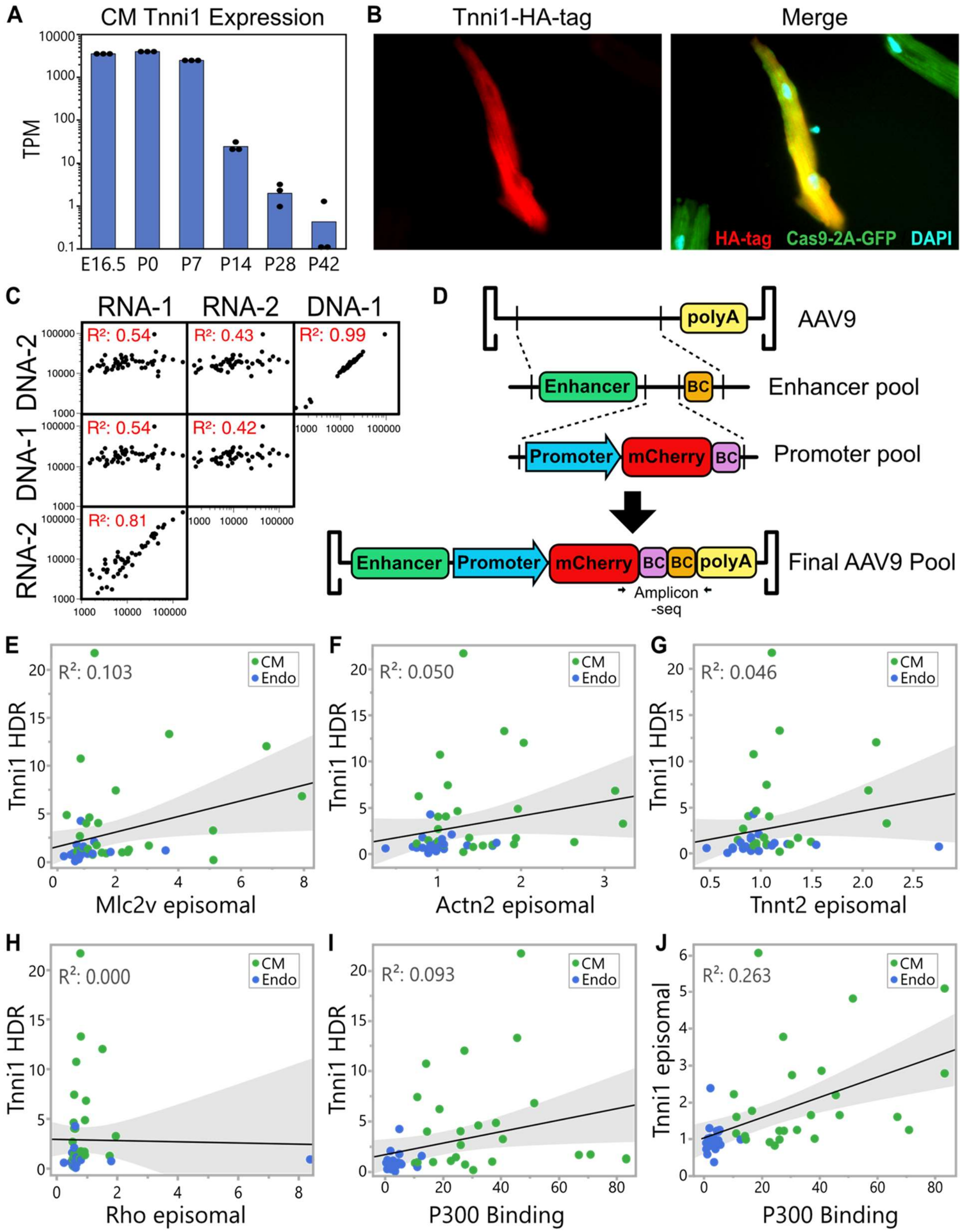


**Supp. Fig. 3. Integrated and episomal MPRAs A.** Tnni1 expression in cardiomyocytes. **B.** HA-tag stained cardiomyocytes from a mouse injected with Tnni1 enhancer-knockin vector. **C.** RNA and DNA samples were collected as duplicates. Sample-to-sample correlations of enhancer frequencies (FPM) are shown. **D.** Cloning strategy for episomal MPRAs. BC, barcode. **E-H.** Comparison of enhancer activities in Tnni1-HDR assay versus an episomal assay with various minimal promoters. **I, J.** Comparison of P300 occupancy at each native enhancer in adult cardiomyocytes to enhancer activities in the Tnni1 HDR assay or episomal assay respectively.

**Supplemental table 1**

Primers used for Sanger sequencing and amplicon-sequencing.

| Primer | Sequence |
|---|---|
| TTN.+AT-F | CCACAGTCACTACTAAGTGCCAT |
| TTN.-T-F | GGACGATGGAGGTGCCAAAA |
| TTN.WT-F | AGTAGCCTGTGCCTTTACTGG |
| TTN-mScarlet-R | ACGCTTCCCAGCCCATTGTT |
| PLN-mScarlet-F | GTTGTACCCCGAGGACGG |
| PLN-Snap-F | CATCGAGGAGTTCCCTGTG |
| PLN-R | ATGCAGATCAGCAGCAGACAT |
| Episomal-MPRA-Round1-F | ACCGGCGGCATGGACGAGCT |
| Episomal-MPRA-Round1-R | CTGGAGTTCAGACGTGTGCTCTTCCGATCTAAGTCAGATGCTCAAGGGGC |
| Episomal-MPRA-Round2-F | AATGATACGGCGACCACCGAGATCTACACTCTTTCCCTACACGACGCTCTTCCGATCTGCATGGACGAGCTGTACAAG |
| Episomal-MPRA-Round2-R (N = index) | CAAGCAGAAGACGGCATACGAGATNNNNNNGTGACTGGAGTTCAGACGT |
| Integrated-MPRA-RNA-Round1-F | ATCGATAATCGGCCGGCCGTC |
| Integrated-MPRA-RNA-Round1-R | CTGGAGTTCAGACGTGTGCTCTTCCGATCTTCCTCAGCCTAGGGCCCCTT |
| Integrated-MPRA-RNA-Round2-F | AATGATACGGCGACCACCGAGATCTACACTCTTTCCCTACACGACGCTCTTCCGATCTTCGGCCGGCCGTCAGCGATC |
| Integrated-MPRA-RNA-Round2-R (N = index) | CAAGCAGAAGACGGCATACGAGATNNNNNNGTGACTGGAGTTCAGACGT |
| Integrated-MPRA-DNA-Round1-F | GTTCAAGCGTCCTCCCCTGC |
| Integrated-MPRA-DNA-Round1-R | CTGGAGTTCAGACGTGTGCTCTTCCGATCTTAGGGTCCCCCCAGTTTCCA |
| Integrated-MPRA-DNA-Round2-F | AATGATACGGCGACCACCGAGATCTACACTCTTTCCCTACACGACGCTCTTCCGATCTTCGGCCGGCCGTCAGCGATC |
| Integrated-MPRA-DNA-Round2-R (N = index) | CAAGCAGAAGACGGCATACGAGATNNNNNNGTGACTGGAGTTCAGACGT |

**Supplemental table 2**

Guide RNA sequences used in this study.

| | **Guide RNA target** | **Sequence** |
|---|---|---|
| sgRNA sequences | TTN.+AT | AGAGAAAAGGGTCTGACCAG |
| | TTN.-T | GGAAGTTACAACTATCACCA |
| | TTN.WT | CGTTCTATGTAAGAGGGCCA |
| | PLN | GGGAGACAAGTTTTTCTAAG |
| | TNNI1 | CTCTAAGCACCTCTACTGGG |

**Supplemental table 3**

Antibodies and stains used in this study.

| | **Antibody (species)** | **Product Number, Manufacturer** |
|---|---|---|
| Primary antibody | HA-Tag (Rb) | C29F4, Cell Signaling |
| | Caveolin 3 (Rb) | PA1066, Life Technologies |
| | Phospholamban (Rb) | 14562s, Cell Signaling |
| | Phospho-Phospholamban (Rb) | 8496s, Cell Signaling |
| | GAPDH (Ms) | MA5-15738, Thermo Fisher |
| Secondary antibody or dye | Gt-anti-Rb Alexa Fluor® Plus 555 | A32732, Thermo Fisher |
| | Gt-anti-Rb Alexa Fluor® Plus 647 | A32733, Thermo Fisher |
| | Gt-anti-Ms Alexa Fluor® Plus 647 | A32728, Thermo Fisher |
| | Anti-Rabbit HRP | 042-206, Protein Simple |
| | SNAP-Cell® 647-SiR | S9102S, NEB |
| | Hoechst | 62249, Thermo Fisher |
| | DAPI | D1306, Thermo Fisher |
| | WGA Alexa Flour® 647 | W32466, Thermo Fisher |

**Supplemental table 4. MPRA enhancers**

Validated enhancers were selected for the MPRA from a search of the VISTA Enhancer Database and the literature (PubMed ID indicated). The cell type the enhancer was reported to be active in is indicated in column A. Enhancers were individually synthesized with a barcode, which was used for the episomal MPRA.

| Enhancer Type | ID | Barcode | Reference coordinates (mm10) | Assayed coordinates (mm10) | Nearest Gene | Reference | Notes |
|---|---|---|---|---|---|---|---|
| Myocardium | 1 | CAGCTTTTGG | chr5:122092218-122094768 | chr5:122092834-122093233 | Myl2 | Vista Enhancer Database #mm77 | - |
| Myocardium | 2 | TGCAGGTTCG | chr18:75502363-75503667 | chr18: 75502708-75503107 | Ctif | Vista Enhancer Database #mm257 | - |
| Myocardium | 3 | GTGGAGTACT | chr19:53715569-53717446 | chr19:53716451-53716850 | Rbm20 | Vista Enhancer Database #mm64 | - |
| Myocardium | 4 | TTTCCGCCGG | chr11:65208855-65210457 | chr11:65209431-65209830 | Myocd | Vista Enhancer Database #mm67 | - |
| Myocardium | 5 | TGGCCGCTGA | chr9:64044784-64046949 | chr9:64045149-64045548 | 1700055C04Rik | Vista Enhancer Database #mm69 | - |
| Myocardium | 6 | CTACAGACTC | chr8:33903821-33907069 | chr8:33905770-33906169 | Rbpms | Vista Enhancer Database #mm70 | - |
| Myocardium | 7 | GTGTACGTTC | chr8:95426508-95428314 | chr8:95426749-95427148 | Cfap20 | Vista Enhancer Database #mm71 | - |
| Myocardium | 8 | ACCAACTTGG | chr3:152303123-152305850 | chr3:152304424-152304823 | Fam73a | Vista Enhancer Database #mm72 | - |
| Myocardium | 9 | TCAAGGTTTG | chr12:83072395-83074413 | chr12:83073211-83073610 | Rgs6 | Vista Enhancer Database #mm78 | - |
| Myocardium | 10 | ATCCGCACCA | chr17:73022163-73024151 | chr17:73022583-73022982 | Lclat1 | Vista Enhancer Database #mm82 | - |
| Myocardium | 11 | CTAGTTAGGA | chr11:65648906-65651643 | chr11:65649932-65650331 | Map2k4 | Vista Enhancer Database #mm85 | - |
| Myocardium | 12 | GATGGGTGAA | chr15:66824301-66827162 | chr15:66826151-66826550 | Tg | Vista Enhancer Database #mm104 | - |
| Myocardium | 13 | AAAGATGGAC | chr12:69757409-69759262 | chr12:69757719-69758118 | Cdkl1 | Vista Enhancer Database #mm122 | - |
| Myocardium | 14 | GTTAGTAACG | chr13:46571320-46574349 | chr13:46571988-46572387 | Cap2 | Vista Enhancer Database #mm132 | - |
| Myocardium | 15 | CTGGGTAGTC | chr1:184041270-184042762 | chr1:184041892-184042291 | Dusp10 | Vista Enhancer Database #mm134 | - |
| Myocardium | 16 | GGGGAGATCT | chr3:104596935-104600543 | chr3:104599431-104599830 | Slc16a1 | Vista Enhancer Database #mm143 | - |
| Myocardium | 17 | TACTCTTAAA | chr4:120391603-120393946 | chr4:120392787-120393186 | Scmh1 | Vista Enhancer Database #mm145 | - |
| Myocardium | 18 | AGGGTTGTCA | chr11:77422469-77424441 | chr11:77423506-77423905 | Ssh2 | Vista Enhancer Database #mm146 | - |
| Myocardium | 19 | GGTTGAACGA | chr18:80330119-80332361 | chr18:80331066-80331465 | Kcng2 | Vista Enhancer Database #mm152 | - |
| Myocardium | 20 | TCAAGCATGA | chr5:21329154-21331111 | chr5:21329618-21330017 | Ccdc146 | Vista Enhancer Database #mm186 | - |
| Myocardium | 21 | CACACGTCAA | chr17:48436330-48438535 | chr17:48436592-48436991 | Apobec2 | Vista Enhancer Database #mm243 | - |
| Myocardium | 22 | AATCTTACTT | chr14:63367789-63369845 | chr14:63368465-63368864 | Blk | Vista Enhancer Database #mm245 | - |
| Myocardium | 23 | CAAGACCCAC | chr12:85454400-85456289 | chr12:85455103-85455502 | Fos | Vista Enhancer Database #mm260 | - |
| Myocardium | 24 | GCATAAAAAA | chr7:110162809-110166321 | chr7:110164032-110164431 | Wee1 | Vista Enhancer Database #mm725 | - |
| Myocardium | 25 | TAACATCGAG | chr4:95317529-95318734 | chr4:95317822-95318221 | Fggy | Vista Enhancer Database #mm308 | - |
| Endocardium | 26 | TCGCAAAGGG | chr2:26571520-26572509 | chr2:26571854-26572253 | Egfl7_E1 | PMID28121289 | - |
| Endocardium | 27 | GCGAATGATC | chr1:133491869-133494057 | chr1:133493441-133493840 | Sox13 | Vista Enhancer Database #mm573 | - |
| Endocardium | 28 | CGCCAATGCA | chr12:79795794-79798372 | chr12:79796258-79796657 | Rad51b | Vista Enhancer Database #mm1393 | - |
| Endocardium | 29 | TCACGTCCTA | chr2:124554786-124555549 | chr2:124554995-124555394 | Sema6d | PMID28121289 | - |
| Endocardium | 30 | GCAGGTCTGG | chr2:26578160-26579122 | chr2:26578438-26578837 | Egfl7_E3 | PMID28121289 | - |
| Endocardium | 31 | TAGCCCCCGT | chr4:139597441-139601336 | chr4:139599079-139599478 | Iffo2 | Vista Enhancer Database #mm1506 | Failed synthesis - not included |
| Endocardium | 32 | TTTCCTCAAA | chr2:26474735-26475664 | chr2:26475026-26475425 | Notch1_E1 | PMID28121289 | - |
| Endocardium | 33 | GAGCTCCCAG | chrX:48005714-48006741 | chrX:48006092-48006491 | Apln | PMID28121289 | - |
| Endocardium | 34 | GAACCCTTAA | chr6:84048677-84049910 | chr6:84048857-84049256 | Dysf | Vista Enhancer Database #hs2170 | Liftover from hg38 chr2:71490876-71494083 |
| Endocardium | 35 | TGACCAAATT | chr4:135253467-135256150 | chr4:135254797-135255196 | Clic4 | Vista Enhancer Database #mm1648 | - |
| Endocardium | 36 | TACTTGAAGT | chr9:37399047-37402046 | chr9:37400570-37400969 | Robo4 | PMID17495228 | - |
| Endocardium | 37 | ATGCGTTGAA | chr4:94745331-94746949 | chr4:94746449-94746848 | Tek | PMID9096345 | - |
| Endocardium | 38 | AGCCGTGCCG | chr11:49632161-49633018 | chr11:49632269-49632668 | Flt4 | PMID19070576 | - |
| Endocardium | 39 | TACATTACCA | chr8:128386820-128387840 | chr8:128387203-128387602 | Nrp1 | PMID19070576 | - |
| Endocardium | 40 | ATCGTGTTAT | chr4:137919926-137920531 | chr4:137920023-137920422 | Ece1 | PMID19070576 | - |
| Endocardium | 41 | AACATCTACC | chr4:115071093-115076331 | chr4:115075532-115075931 | Tal1 | PMID14966269 | - |
| Endocardium | 42 | AGTGAATAGC | chr2:119327125-119327925 | chr2:119327153-119327552 | Dll4 | PMID23830865 | - |
| Endocardium | 43 | CGGACAGTCG | chr6:88202604-88203797 | chr6:88202958-88203357 | Gata2 | PMID17395646 | - |
| Endocardium | 44 | CAGGAGAATG | chr13:83571928-83572369 | chr13:83571999-83572398 | Mef2c | PMID15501228 | - |
| Endocardium | 45 | AAGTTATGCT | chr17:34564510-34565332 | chr17:34564815-34565214 | Notch4 | PMID15684396 | - |
| Endocardium | 46 | TGGGTTCTAG | chr19:37435671-37435904 | chr19:37435625-37436024 | Hhex | PMID15649946 | - |
| Endocardium | 47 | TGGCAACTCA | chr5:75961867-75962690 | chr5:75962135-75962534 | Kdr | PMID27079877 | - |
| Endocardium | 48 | TCTTACAGCG | chr18:13847776-13849408 | chr18:13848506-13848905 | Zfp521 | Vista Enhancer Database #hs1653 | Liftover from hg38 chr18:25228374-25230020 |
| Endocardium | 49 | GGGGCACGCG | chr4:57845561-57847767 | chr4:57846984-57847383 | Pakap | Vista Enhancer Database #mm261 | - |
| Endocardium | 50 | GGAGTATAAA | chr19:37492022-37497349 | chr19:37493919-37494318 | 1700122C19Rik | Vista Enhancer Database #hs1866 | Liftover from hg38 chr10:92754239-92758228 |
| ESC | E1 | GTGGTATGTG | N/A | chr2:4512536-4512936 | Prpf18 | PMID28121289 | - |
| ESC | E2 | GTTCTTCTGC | N/A | chr13:6040082-6040482 | Klf6 | PMID28121289 | - |
| ESC | E3 | ATTCTCTGCA | N/A | chr11:5046317-5046717 | Rasl10a | PMID28121289 | - |
| ESC | E4 | GCTATGTAGG | N/A | chr7:4902457-4902857 | Zfp628 | PMID28121289 | - |
| ESC | E5 | AGACGCTAGA | N/A | chr8:3421492-3421892 | Arhgef18 | PMID28121289 | - |

**Supplemental Table 5: MPRA promoters**

Short promoter sequences overlapping the TSS of the indicated genes were selected for this study. Additional short minimal promoters were selected from the literature.

| Promoter Name | Species | Sequence (5' --> 3') | Barcode | Source |
|---|---|---|---|---|
| Tnnt2 | Mouse | TCTGAGCAGCTGGAGGACCACATGAGCTTATATGGCGTGGGGTACTTGTTCTTTTAGCCCTGTGCCGG<br>GCACCTGCCAAAATAGCAGCCAACACCCCCCATTGTGTTGTTCCCCCCCCCCCCCATCTCCTGCTGCAC<br>ATTCCTCCCTCCGCGGGGCTTGGCTCACAAGGCCCCAGCCCACATGCCTGCTTAAAGCTCTCC | TACGGCCTAG | this study |
| TnnI1 | Mouse | GCAGCACATATCTGCCCTGCGAGGTGTGCAGGGTGGGAGGGGGTGGGAAGGAGGGGCAGCTGGAG<br>GGGCAGCGGCTGTTCTATTTTTACTGGCCAGTTGCCGGAGGCCACGGTTTTCATAGCCTGCCCTCAGC<br>TCTGCCCCCACACTCTGCAGTCTGTGGTGAGGCTCAGGCCAGCCTAGCTCCACGAGGACTAAACTAG | TCCTAAATAA | this study |
| Actn2 | Mouse | GGAACCTGAGCGGGGCTCTCCAGCCAGCCAACCACACGTGTTGGAGACGGGGAGTGGGTGGGGCCG<br>GCAGCACGTGACTCTCAGCGGGCTATTAATCCGCGCGGCGCTGCCTGCAGGCGTGCTGGTACTTCGC<br>CGGAGACTCCGCGCCCAGGCGTGCCGCCCCAGGAACCGCGAGAAGGTCACCGCAGCCCGCCGCCTTT | TATTCACGAA | this study |
| Rho | Cow | GATTCAGCCGGGAGCTTAGGGAGGGGAGGTCACTTCATAAGGGCTTGGGGGGGGAGTTGGAGCCA<br>CGAGTCGTCCAGCCGGAGCCCCGTGTGGCTGAGCTCCGGCCTCAGAAGCATCCCCGGGATCCACCGG<br>TCGCCA | ATACACTGCG | PMID: 26576614 |
| Mlc2v | Rat | AGACAATGGCAGGACCCAGAGCACAGAGCATCGTTCCCAGGCCAGGCCCCAGCCACTGTCTCTTTAA<br>CCTTGAAGGCATTTTTGGGTCTCACGTGTCCACCCAGGCGGGTGTCGGACTTTGAACGGCTCTTACTT<br>CAGAAGAACGGCATGGGGTGGGGGGGCTTAGGTGGCCTCTGCCTCACCTACAACTGCCAAAAGTGG<br>TCATGGGGTTATTTTTAACCCCAGGGAAGAGGTATTTATTGTTCCACAGCAGGGGCCGGCCAGCAGG<br>CTCCTTGAATT | TAAATACTGT | PMID: 2808370 |

**Supplemental table 6 - Activity for all enhancer-promoter pairs**

Activity was calculated as the ratio of frequency in RNA/DNA, and normalized such that the average activity of the endothelial enhancers for each promoter was 1.0. Column "Tnni1UTR" acitivies are from the integrated MPRA, and "Mlc2v" to "Rho" are from the episomal MPRA. "P300 occupancy" reflects cardiomyocyte specific P300 occupancy at each enhancer region at P28.

| Enhancer ID | mm10 coordinates | Enhancer Type | Tnni1UTR | Mlc2v | Tnni1 | Actn2 | TnT | Rho | P300 occupancy |
|---|---|---|---|---|---|---|---|---|---|
| 1 | chr5:122092218-122094768 | Myocardial | 4.880811 | 0.44483 | 1.015748 | 1.96218 | - | 0.933711 | 38.2306 |
| 2 | chr18:75502363-75503667 | Myocardial | 4.629611 | 1.169266 | 1.251969 | 1.238042 | 0.951902 | 0.560226 | 32.1357 |
| 3 | chr19:53715569-53717446 | Myocardial | 1.281334 | 2.427499 | 2.787402 | 2.6396 | 1.49217 | 1.748585 | 83.2251 |
| 4 | chr11:65208855-65210457 | Myocardial | 1.748101 | 1.398036 | 1.251969 | 1.378198 | 1.029083 | 0.797898 | 70.9049 |
| 5 | chr9:64044784-64046949 | Myocardial | 0.897714 | 1.703062 | 2.220472 | 1.541713 | 0.926174 | 0.780922 | 10.2621 |
| 6 | chr8:33903821-33907069 | Myocardial | 1.359469 | 1.118429 | 5.102362 | 1.004449 | - | 0.933711 | 83.2194 |
| 7 | chr8:95426508-95428314 | Myocardial | 4.053203 | 1.448873 | 1.228346 | 1.097887 | 0.87472 | 0.577203 | 26.0068 |
| 8 | chr3:152303123-152305850 | Myocardial | 6.839292 | 7.943385 | 4.818898 | 3.130145 | 2.058166 | 0.967664 | 51.4636 |
| 9 | chr12:83072395-83074413 | Myocardial | 0.753416 | 1.270942 | 1.228346 | 1.424917 | 0.951902 | 0.594179 | 27.5943 |
| 10 | chr17:73022163-73024151 | Myocardial | 1.706236 | 3.062969 | 1.606299 | 1.938821 | 1.363535 | 0.916734 | 66.8625 |
| 11 | chr11:65648906-65651643 | Myocardial | 0.975908 | 1.563258 | 1.606299 | 1.074527 | 0.951902 | 0.526273 | 11.1579 |
| 12 | chr15:66824301-66827162 | Myocardial | 7.441511 | 2.008088 | 1.15748 | 1.121246 | 1.05481 | 0.577203 | 11.0729 |
| 13 | chr12:69757409-69759262 | Myocardial | 10.75035 | 0.889659 | 1.086614 | 1.027809 | 0.926174 | 0.645109 | 14.1118 |
| 14 | chr13:46571320-46574349 | Myocardial | 13.30717 | 3.71115 | 2.19685 | 1.798665 | 1.183445 | 0.797898 | 45.5615 |
| 15 | chr1:184041270-184042762 | Myocardial | 0.205103 | 5.121895 | 2.740157 | 1.30812 | 1.183445 | 0.560226 | 30.3951 |
| 16 | chr3:104596935-104600543 | Myocardial | 21.72777 | 1.334489 | 1.653543 | 1.30812 | 1.106264 | 0.780922 | 46.8825 |
| 17 | chr4:120391603-120393946 | Myocardial | 4.023926 | 1.067591 | 0.992126 | 1.004449 | 1.080537 | 0.611156 | 14.435 |
| 18 | chr11:77422469-77424441 | Myocardial | 12.03856 | 6.812247 | 3.779528 | 2.032258 | 2.135347 | 1.510914 | 27.3448 |
| 19 | chr18:80330119-80332361 | Myocardial | 3.274014 | 5.109185 | 2.858268 | 3.223582 | 2.238255 | 1.952304 | 40.6119 |
| 20 | chr5:21329154-21331111 | Myocardial | 6.249733 | - | 6.070866 | 0.770857 | - | - | 18.7383 |
| 21 | chr17:48436330-48438535 | Myocardial | 2.694906 | 0.86424 | 0.992126 | 1.004449 | 0.823266 | 0.526273 | 26.0692 |
| 22 | chr14:63367789-63369845 | Myocardial | 1.11813 | 0.775274 | 0.92126 | 0.747497 | 0.951902 | 0.611156 | 22.6674 |
| 23 | chr12:85454400-85456289 | Myocardial | 1.042001 | 2.38937 | 1.653543 | 1.915462 | 1.05481 | 0.780922 | 36.9493 |
| 24 | chr7:110162809-110166321 | Myocardial | 1.477818 | 0.940497 | 0.826772 | 0.887653 | 0.771812 | 0.577203 | 24.4079 |
| 25 | chr4:95317529-95318734 | Myocardial | 0.998724 | 2.211438 | 1.771654 | 1.658509 | 1.286353 | 0.763945 | 16.5384 |
| 26 | chr2:26571520-26572509 | Endothelial | 0.848888 | 1.105719 | 1.228346 | 1.027809 | 0.926174 | 0.662086 | 4.39967 |
| 27 | chr1:133491869-133494057 | Endothelial | 4.274455 | 0.889659 | 0.968504 | 0.911012 | 0.926174 | 0.611156 | 4.87592 |
| 28 | chr12:79795794-79798372 | Endothelial | 0.954853 | 0.915078 | - | - | 1.543624 | - | 4.36565 |
| 29 | chr2:124554786-124555549 | Endothelial | 0.517701 | - | - | - | 0.720358 | 0.509297 | 11.0785 |
| 30 | chr2:26578160-26579122 | Endothelial | 0.079795 | 0.711727 | 1.015748 | 0.887653 | 0.668904 | 0.628133 | 3.42448 |
| 32 | chr2:26474735-26475664 | Endothelial | 1.793321 | 0.762565 | 1.204724 | 0.817575 | 0.900447 | 0.560226 | 4.87025 |
| 33 | chrX:48005714-48006741 | Endothelial | 0.519217 | 0.775274 | 1.015748 | 1.074527 | 0.926174 | 0.662086 | 1.27001 |
| 34 | chr6:84048677-84049910 | Endothelial | 0.928653 | - | - | 1.354839 | - | 8.369442 | 0.34018 |
| 35 | chr4:135253467-135256150 | Endothelial | 1.607659 | 0.978625 | 0.992126 | 1.051168 | 0.900447 | 0.509297 | 12.5923 |
| 36 | chr9:37399047-37402046 | Endothelial | 0.301802 | 0.749856 | 0.897638 | 0.887653 | 0.900447 | 0.526273 | 0.680361 |
| 37 | chr4:94745331-94746949 | Endothelial | 0.525943 | 0.724437 | 0.826772 | 0.911012 | 0.823266 | 0.594179 | 2.35858 |
| 38 | chr11:49632161-49633018 | Endothelial | 0.652498 | 0.87695 | 0.944882 | 0.957731 | 0.848993 | 0.628133 | 1.7009 |
| 39 | chr8:128386820-128387840 | Endothelial | 0.657133 | 0.86424 | 0.850394 | 0.817575 | 0.823266 | 0.526273 | 3.56623 |
| 40 | chr4:137919926-137920531 | Endothelial | 1.043112 | 1.842865 | 0.590551 | 1.074527 | 1.260626 | - | 1.08858 |
| 41 | chr4:115071093-115076331 | Endothelial | 1.148077 | 0.902369 | 0.92126 | 0.98109 | 1.131991 | 0.645109 | 2.67609 |
| 42 | chr2:119327125-119327925 | Endothelial | 2.103845 | 1.321779 | 1.299213 | 1.167964 | 0.977629 | 0.577203 | 1.8143 |
| 43 | chr6:88202604-88203797 | Endothelial | 0.75082 | 0.571924 | 2.385827 | - | 2.752796 | 1.799515 | 2.15448 |
| 44 | chr13:83571928-83572369 | Endothelial | 0.869685 | 1.270942 | 1.251969 | 1.658509 | 1.106264 | 0.797898 | 5.31815 |
| 45 | chr17:34564510-34565332 | Endothelial | 0.279387 | 0.826112 | 1.03937 | 0.98109 | 0.977629 | 0.577203 | 2.09778 |
| 46 | chr19:37435671-37435904 | Endothelial | 0.658034 | 0.673599 | 0.732283 | 0.747497 | 0.720358 | 0.475344 | 3.88373 |
| 47 | chr5:75961867-75962690 | Endothelial | 0.773939 | 0.622761 | 0.850394 | 0.700779 | 0.848993 | 0.509297 | 5.60164 |
| 48 | chr18:13847776-13849408 | Endothelial | 0.915348 | 0.66089 | 0.874016 | 0.911012 | 1.05481 | 0.594179 | 2.47765 |
| 49 | chr4:57845561-57847767 | Endothelial | 0.59404 | 0.355864 | 0.377953 | 0.373749 | 0.463087 | 0.237672 | 3.41881 |
| 50 | chr19:37492022-37497349 | Endothelial | 1.201792 | 3.596765 | 0.732283 | 1.705228 | 0.797539 | - | 0.725718 |

**Supplemental Table 7 - Plasmid sequences**

Experiments were conducted in vivo, with constructs delivered via standard recombinant AAV2 genome, packaged in an AAV9 capsid. Plasmid sequences, spanning from viral ITR to ITR, are included here for all experiments.

**AAV-HDR-TTN.WT-mScarlet-Fusion** - inserts mScarlet at C-terminus of wildtype Titin.

CTGCGCGCTCGCTCGCTCACTGAGGCCGCCCGGGCAAAGCCCGGGCGTCGGGCGACCTTTGGTCGCCCGGCCTCAGTGAGCGAGCGAGCGCGCAGAGAGGGAGTGGCCAACTCCATCACTAGGGGTTCCTTGTAGTTAATGATTAACCCGCCATGCTACTTATCTACCAGGGTAATGGGCT
CGAGGAGGGCCTATTTCCCATGATTCCTTCATATTTGCATATACGATACAAGGCTGTTAGAGAGATAATTGGAATTAATTTGACTGTAAACACAAAGATATTAGTACAAAATACGTGACGTAGAAAGTAATAATTTCTTGGGTAGTTTGCAGTTTTAAAATTATGTTTTAAAATGGACTATCATAT
GCTTACCGTAACTTGAAAGTATTTCGATTTCTTGGCTTTATATATCTTGTGGAAAGGACGAAACACCGCGTTCTATGTAAGAGGGCCAGTTTTAGAGCTAGAAATAGCAAGTTAAAATAAGGCTAGTCCGTTATCAACTTGAAAAAGTGGCACCGAGTCGGTGCTTTTTTGGTACCATTGTTCCT
GCAGGCAGATGTCACCGAGTAACGAGATGTCTTTGATGTCCCCCGTGTATCTGCTATGCTCTACAGCTCTGGTGGAAGAACCACCGAGAGAGGTGGTATTGAAAACGAGCAGTGACGTGAGCTTGCACGGAAGCGTCTCGTCTCAGTCAGTCCAAATGTCTGCCTCTAAGCAGGAGGCATCC
TTCAGCAGCTTCAGCAGCAGCAGCGCTAGCAGCATGACTGAAATGAAGTTTGCGAGCATGTCTGCCCAAAGCATGTCTTCCATGCAAGAGTCCTTTGTAGAAATGAGTTCCAGCAGTTTTATGGGAAAATCTAGCATGACACAACTGGAAAGCTCAACCAGTAGAATGCTCAAAGCAGGCGG
AAGAGGTGAGCAAAAGAGTTCCTGGTAGAGGTAGATCAAGCCACCTAGCATGGCATCGTTCTAGAGCAGTGGTTCTCCGCCTTCCTAATGCTGCGACCCTTTAATACAGTTCCTCACGTTGTCGAGACTCTCAATCATGAAATTATTTTTGTTGCTACTTCCCAACTGTAATTTTGCTAACTTTAT
GAGTCGTAATGTAAATATGTTTGGAGGTACGCGTTGGCCACAGGGGTCACGGCCCACATGTTGAGAACCACTGTCTTAGTGCATTAAAAGTCAATAGAAAGCGAATAGGATTTCTTGTTTTCGGTGACCCTAAAACTTTTCTGTTGCTGTTTCATAAAGGAATTCCACCTAAAATTGAAGCTCTT
CCCTCTGATATCAGCATCGACGAAGGCAAAGTCCTAACAGTAGCCTGTGCCTTTACTGGGGAGCCTACTCCAGAAATAACATGGTCCTGTGGTGGAAGGAAAATCCAAAACCAAGAACAACAAGGAAGATTCCACATTGAAAACACAGATGACCTGACCACCCTGATCATCATGGACGTACA
GAAACAAGATGGCGGACTTTATACCCTAAGCTTAGGGAATGAATTTGGATCTGACTCTGCCACTGTGAATATTAATATCCGTTCTATGATGGTGAGCAAGGGCGAGGCAGTGATCAAGGAGTTCATGCGGTTCAAGGTGCACATGGAGGGCTCCATGAACGGCCACGAGTTCGAGATCGAG
GGCGAGGGCGAGGGaCGCCCCTACGAGGGCACCCAGACCGCCAAGCTGAAGGTGACCAAGGGTGGCCCCCTGCCCTTCTCCTGGGACATCCTGTCCCCTCAGTTCATGTACGGCTCCAGGGCCTTCACCAAGCACCCCGCCGACATCCCCGACTACTATAAGCAGTCCTTCCCCGAGGGCTT
CAAGTGGGAGCGCGTGATGAACTTCGAGGACGGCGGCGCCGTGACCGTGACCCAGGACACCTCCCTGGAGGACGGCACCCTGATCTACAAGGTGAAGCTCCGCGGCACCAACTTCCCTCCTGACGGCCCCGTAATGCAGAAGAAaACAATGGGCTGGGAAGCGTCCACCGAGCGGTTGTA
CCCCGAGGACGGCGTGCTGAAGGGCGACATTAAGATGGCCCTGCGCCTGAAGGACGGCGGaCGCTACCTGGCGGACTTCAAGACCACCTACAAGGCCAAGAAGCCCGTGCAGATGCCCGGCGCCTACAACGTCGACCGCAAGTTGGACATCACCTCCCACAACGAGGACTACACCGTGGT
GGAACAGTACGAACGCTCCGAGGGtCGCCACTCCACCGGtGGtATGGACGAGCTGTACAAGTAGGAGGGCCAGGGCCAAGAGCGGGGGCCCTTTACACTCGATGCTCATTCATAATCTCTCTCAAATGATTCACATGAACTAATCTTTCTGACCTGTAAATATTTTTAAGTAGTTTTATATCCAT
CTGAATGAGTCACAGTTCCAAAACTATGCATTTCAATCCTTTCATACTTGACCTGAGAATATTTGCTAATGACACACATGTACATACTGTATATAGCCGGGTTCACGGTTATAAAGTTTTGTACCATTTTATTTTATGACATTTTACACTGTAACTTTTGAAACTAACTGTTGGTGGGAGAAAGTTTC
TTATAGATTATAGAACGAATACCCTGCCAATACTTAATCAATCTTTGTGCCTCAACATACTGTTGATGTCTAAGTATGCCTCAGTGGGTTGAGAAATTCCCCACTGAAGATGTCCTATCCACCTAAAAAAAAAAGGATGCTTCTGTACATGTCCCTGACATGTGCACTGACACCTATTGAATCAGA
AATGTAAGGCATTGGTGATGTTTGCATCTACCCTCCTGTAAGCAACACTTTAACGTCTTACGTTTTCTCTGATGATGTCACACTGAAAATTATCATGACAAATATAACCAGAGCAAAGTGTAACGGCCATACACTTTGTTCGCCCACTTCACACTGTCTCTGGCATACAGAGTGCATGGATAGCTT
GGGAAATGTACCATCTCCTGGCCATCCCCTCGTCTAACCAAGCTATGCAAGTATTCCTATGCCAGAGCGGTGTCGACTCCTGGAGGTCCCAGGTGCAGCTCCTGCCTTGTGTGGTAGTTCTAAATTCAAATTTCACCGGGAACTGGGCAACTAAGCAATGAGCCACAGCAAAATAATAATAATA
ATAATTTAAAAAAATGTAACAAGTGAAGCTGTCGTTAAAACAACAATACTACCAACTGCCCATAAAAAAAAAAAATTACTAATTTGATGTAGTCCTTTTCATGCCAGTATATGTTCAATTGGTAGTTATAAATGTTGCGATCGCACATGGGCTAGCGGACTTTGTACACTGGATACTAGTCCCATTA
CCCTGGTAGATAAGTAGCATGGCGGGTTAATCATTAACTACAAGGAACCCCTAGTGATGGAGTTGGCCACTCCCTCTCTGCGCGCTCGCTCGCTCACTGAGGCCGGGCGACCAAAGGTCGCCCGACGCCCGGGCTTTGCCCGGGCGGCCTCAGTGAGCGAGCGAGCGCGCAG

**AAV-HDR-TTN.AT63117fs-mScarlet** - inserts mScarlet after truncation caused by AT63117 mutation.

CTGCGCGCTCGCTCGCTCACTGAGGCCGCCCGGGCAAAGCCCGGGCGTCGGGCGACCTTTGGTCGCCCGGCCTCAGTGAGCGAGCGAGCGCGCAGAGAGGGAGTGGCCAACTCCATCACTAGGGGTTCCTTGTAGTTAATGATTAACCCGCCATGCTACTTATCTACCAGGGTAATGGGCT
CGAGGAGGGCCTATTTCCCATGATTCCTTCATATTTGCATATACGATACAAGGCTGTTAGAGAGATAATTGGAATTAATTTGACTGTAAACACAAAGATATTAGTACAAAATACGTGACGTAGAAAGTAATAATTTCTTGGGTAGTTTGCAGTTTTAAAATTATGTTTTAAAATGGACTATCATAT
GCTTACCGTAACTTGAAAGTATTTCGATTTCTTGGCTTTATATATCTTGTGGAAAGGACGAAACACCGAGAGAAAAGGGTCTGACCAGGTTTTAGAGCTAGAAATAGCAAGTTAAAATAAGGCTAGTCCGTTATCAACTTGAAAAAGTGGCACCGAGTCGGTGCTTTTTTGGTACCATTGTTCC
TGCAGGCTCTTGTTCAGATCCCCCAGGACCACCTTCAAATGCACATGTCACTGATACCACCAAGAAATCGGCTTCTTTAGCATGGGGCAAGCCTCATTATGATGGTGGGCTTGAAATCACTGGCTATGTGGTAGAGCATCAAAAAGTAGGAGATGATGCCTGGATCAAAGACACAACAGGAAC
AGCCCTCAGAATCACTCAGTTTGTGGTTCCTGATCTTCAGACTAAGGAAAAATACAATTTTAGAATTAGTGCCATCAACGATGCAGGTGTGGGGGAACCAGCTGTGATTCCAAATGTTGAAATTGTAGAGAAGGAAGTGGCTCCTGATTTTGAGCTAGATGCTGAGCTGCGAAGAACACTTGT
GGTTAGGGCAGGGCTCAGTATTAGGATTTTTGTGCCAATTAAAGGTCGTCCTGCACCTGAAGTGACATGGACCAAAGATAATATCAACCTGAAGCACCGAGCCAACATCGAAAACACAGAGTCATTTACTCTTCTCATTATTCCAGAATGTAACAGATATGACACTGGTAAATTTGTGATGACC
ATCGAAAACCCAGCTGGGAAGAAAAGCGGCTTCGTTAACGTCAGAGTCCTAGACACACCAGGCCCTGTCCTTAACCTAAGGCCTACAGACATCACAAAGGACAGTGTCACCCTACACTGGGACCTACCTCTGATAGATGGGGGCTCACGTATAACAAATTATATTGTGGAGAAACGTGAAGC
AACCAGGAAATCCTACTCCACAGTCACTACTAAGTGCCATAAATGTACATATAAAGTGACTGGCTTGACAGAAGGCTGTGAATATTTCTTCAGAGTGATGGCAGAAAATGAATATGGGGTTGGTGAGCCAACGGAAACCACAGAACCTGTGAGAGCCTCTGAGGCACCGTTACCCCCAGATA
GCCTTAACATTATGGACATTACCAAGAACACAGTCAGCCTGGCGTGGCCTAAGCCAAGGCATGATGGTGGGAGCAAGATATCACTGGCTATGGCGGCCGCcATGGTGAGCAAGGGCGAGGCAGTGATCAAGGAGTTCATGCGGTTCAAGGTGCACATGGAGGGCTCCATGAACGGCCACG
AGTTCGAGATCGAGGGCGAGGGCGAGGGaCGCCCCTACGAGGGCACCCAGACCGCCAAGCTGAAGGTGACCAAGGGTGGCCCCCTGCCCTTCTCCTGGGACATCCTGTCCCCTCAGTTCATGTACGGCTCCAGGGCCTTCACCAAGCACCCCGCCGACATCCCCGACTACTATAAGCAGTCC
TTCCCCGAGGGCTTCAAGTGGGAGCGCGTGATGAACTTCGAGGACGGCGGCGCCGTGACCGTGACCCAGGACACCTCCCTGGAGGACGGCACCCTGATCTACAAGGTGAAGCTCCGCGGCACCAACTTCCCTCCTGACGGCCCCGTAATGCAGAAGAAaACAATGGGCTGGGAAGCGTCC
ACCGAGCGGTTGTACCCCGAGGACGGCGTGCTGAAGGGCGACATTAAGATGGCCCTGCGCCTGAAGGACGGCGGaCGCTACCTGGCGGACTTCAAGACCACCTACAAGGCCAAGAAGCCCGTGCAGATGCCCGGCGCCTACAACGTCGACCGCAAGTTGGACATCACCTCCCACAACGAG
GACTACACCGTGGTGGAACAGTACGAACGCTCCGAGGGtCGCCACTCCACCGGtGGtATGGACGAGCTGTACAAGTAGGGCGCGCCCCAGTGGACCCACATCTCAACAGTGAAAGGGTTAGAATGTGTGGTGAGGAACCTAACCGAAGGAGAGGAATATACCTTCCAAGTGATGGCGGTA
AACAGTGCAGGCAGGAGTGCCCCTAGAGAGAGCAGACCTGTCATTGTCAAAGAGCAGACGATGCTCCCAGAGCTGGATCTCCGTGGCATCTATCAGAAGCTAGTCATTGCCAGAGCTGGTGACAACATCAAAGTTGAAATCCCAGTACTTGGTCGACCCAAGCCCACAGTCACATGGAAAA
AGGGAGACCAAATCCTTAAACAGACACAGAGAGTTAATGTTGAAAACACAGCAACGTCAACCATCTTAAATATCAATGAGTGTGTCAGAAGTGACAGTGGACCCTATCCATTAACAGCAAAGAACACTGTAGGAGAGGTTGGTGACGTTATAACCATTCAAGTCCATGATATTCCAGGACCA
CCTACTGGACCCATTAAATTTGATGAAGTTTCATCTGACTTTGTAACCTTCTCTTGGGAGCCACCAGAGAATGATGGAGGTGTACCAATAAGCAACTATGTAGTAGAGATGAGACAGACAGACAGTACTACATGGGTGGAGTTAGCAACAACTGTTATACGTACTACCTATAAAGCAACTCGTC
TGACCACAGGAGTGGAGTATCAATTCCGTGTAAGGGCTCAAAACAGATATGGAGTTGGACCGGGCATCACATCTGCATCTGTAGTTGCAAACTATCCATTCAAGGTGCCCGGGCCTCCCGGTACCCCTCAGGTTACTGCAGTCACCAAAGACTCCATGACAATCAGCTGGCATGAACCACTTT
CTGATGGTGGAAGCCCCATTTTAGGATATCACATTGAAAGGAAAGAACGGAATGGCATCCTCTGGCAGACTGTGAGCAAAGCATTAGTACCAGGCAACATTTTCAAATCAACTGGACTTACCGATGGCATTGCTTATGAATTCCGAGTAATTGCAGAAAACATGGCAGGCAAGAGCGCGATC
GCACATGGGCTAGCGGACTTTGTACACTGGATACTAGTCCCATTACCCTGGTAGATAAGTAGCATGGCGGGTTAATCATTAACTACAAGGAACCCCTAGTGATGGAGTTGGCCACTCCCTCTCTGCGCGCTCGCTCGCTCACTGAGGCCGGGCGACCAAAGGTCGCCCGACGCCCGGGCTTTG
CCCGGGCGGCCTCAGTGAGCGAGCGAGCGCGCAG

**AAV-HDR-TTN.P22582fs-mScarlet** - inserts mScarlet after truncation caused by P22582fs mutation.

CTGCGCGCTCGCTCGCTCACTGAGGCCGCCCGGGCAAAGCCCGGGCGTCGGGCGACCTTTGGTCGCCCGGCCTCAGTGAGCGAGCGAGCGCGCAGAGAGGGAGTGGCCAACTCCATCACTAGGGGTTCCTTGTAGTTAATGATTAACCCGCCATGCTACTTATCTACCAGGGTAATGGGCT
CGAGGAGGGCCTATTTCCCATGATTCCTTCATATTTGCATATACGATACAAGGCTGTTAGAGAGATAATTGGAATTAATTTGACTGTAAACACAAAGATATTAGTACAAAATACGTGACGTAGAAAGTAATAATTTCTTGGGTAGTTTGCAGTTTTAAAATTATGTTTTAAAATGGACTATCATAT
GCTTACCGTAACTTGAAAGTATTTCGATTTCTTGGCTTTATATATCTTGTGGAAAGGACGAAACACCGGAAGTTACAACTATCACCAGTTTTAGAGCTAGAAATAGCAAGTTAAAATAAGGCTAGTCCGTTATCAACTTGAAAAAGTGGCACCGAGTCGGTGCTTTTTTGGTACCATTGTTCCTG
CAGGTGAATTCCGAGTAATTGCAGAAAACATGGCAGGCAAGAGCAAGCCAAGCAAGCCATCTGAACCAATGTTTGCTTTGGATCCCATTGACCCACCCGGGAAACCAGTTCCTCTAAACATTACAAGACACACAGTGGCACTGAAGTGGGCTAAGCCTGAATATACCGGAGGCTTTAAAATT
ACTAGCTATGTGGTTGAAAAGAGGGACCTTCCTAATGGACGATGGCTGAAGGCCAACTTCAGCAACATTTTAGAGAATGAATTTACGGTCAGCGGCCTAACTGAAGATGCTGCATACGAGTTCCGTGTGATTGCCAAAAATGCTGCCGGTGCCATCAGTCCACCGTCTGAGCCATCAGATGCC
ATCACATGCAGGGACGACCTGGAGGCACCAAGAATAATGGTGGATGTTAGATTTAAGGATACCATCACATTAAAAGCTGGCGAAGCATTCAAGTTGGAAGCTGATGTTTCAGGTCGCCCACCTCCAACAATGGAATGGACTAAAGATGGAAAGGAGCTTGAAGGCACAGGGAAATTAGAA
ATAAAAATTGCAGACTTCTCCACTCATCTGATAAACAAGGATTCCTCAAGAACGGATAGTGGGGCTTACATCCTGACAGCAACCAATCCTGGTGGCTTTGCCAAACACATTTTCAATGTCAAAGTTCTTGATAGACCGGGACCACCTGAGGGACCTTTGGCTGTGTCTGATGTGACATCAGAGA
AGTGTGTGTTGTCATGGTTGCCTCCACTGGACGATGGAGGTGCCAAAATTGATCATTACATCGTACAGAAACGTGAAACCAGCAGGTTGGCATGGACAAATGTTGCCACAGAAGTCCAAGTAACAAAGCTGAAGGTTACTAAGCTCCTGAAAGGGAATGAATACATATTCCGTGTCATGGCA
GTAAATAAATACGGGGTTGGAGAGCCACTTGAATCAGAACCTGTGCTTGCAGTGGATCCATATGGACCACCGACCCACCCAAGAACCCGGAAGATGGTGAGCAAGGGCGAGGCAGTGATCAAGGAGTTCATGCGGTTCAAGGTGCACATGGAGGGCTCCATGAACGGCCACGAGTTCGAG
ATCGAGGGCGAGGGCGAGGGaCGCCCCTACGAGGGCACCCAGACCGCCAAGCTGAAGGTGACCAAGGGTGGCCCCCTGCCCTTCTCCTGGGACATCCTGTCCCCTCAGTTCATGTACGGCTCCAGGGCCTTCACCAAGCACCCCGCCGACATCCCCGACTACTATAAGCAGTCCTTCCCCGA
GGGCTTCAAGTGGGAGCGCGTGATGAACTTCGAGGACGGCGGCGCCGTGACCGTGACCCAGGACACCTCCCTGGAGGACGGCACCCTGATCTACAAGGTGAAGCTCCGCGGCACCAACTTCCCTCCTGACGGCCCCGTAATGCAGAAGAAaACAATGGGCTGGGAAGCGTCCACCGAGCG
GTTGTACCCCGAGGACGGCGTGCTGAAGGGCGACATTAAGATGGCCCTGCGCCTGAAGGACGGCGGaCGCTACCTGGCGGACTTCAAGACCACCTACAAGGCCAAGAAGCCCGTGCAGATGCCCGGCGCCTACAACGTCGACCGCAAGTTGGACATCACCTCCCACAACGAGGACTACAC
CGTGGTGGAACAGTACGAACGCTCCGAGGGtCGCCACTCCACCGGtGGtATGGACGAGCTGTACAAGTAGCTATCACCAAGGACTCCATGGTCGTCTGCTGGGGGCATCCTGACTCTGATGGTGGAAGTGAAATTATCAACTACATTGTGGAGCGGCGTGATAAAGCTGGCCAGCGCTGGGT
AAAATGCAACAAAAAAGCTCTGACTGACTTAAGATTTAAAGTATCTGGGCTGACAGAAGGACATGAATACGAGTTCAGAATTATGGCAGAAAACGCAGCTGGAATCAGTGCTCCGAGTGCTACCAGCCCGTTTTATAAAGCTTGCGACACTGTGTTTAAACCTGGCCCACCGGGTAACCCAC
GTGTCCTTGATACAAGCAGGTCGTCCATTTCAATTGCGTGGAATAAACCCATTTATGATGGAGGTTCAGAAATCACTGGGTATATGGTTGAGATTGCCCTGCCTGAAGAAGATGAATGGCAGGTTGTCACTCCACCAGCTGGGCTCAAGGCAACTTCCTATACCATCACCAGCCTCATAGAGA
ATCAGGAATATAAAATCCGCATTTATGCCATGAATTCTGAGGGACTTGGGGAACCTGCCCTTGTTCCCGGAACTCCAAAGGCAGAAGAAAGAATGTTGCCTCCAGAGATAGAACTGGATGCTGACTTACGCAAAGTTGTCACTATACGAGCCTGCTGCACCTTGAGGCTCTTTGTTCCCATCAA
AGGGAGACCAGCACCTGAGGTCAAGTGGGCCCGAGAGCATGGAGAATCTCTGGACAAGGCTAGCATTGAATCCACAAGCTCTTACACCCTGCTTGTTGTTGGAAATGTTAACAGATTCGACAGTGGCAAGTACATACTAACGGTAGAAAACAGCTCTGGCAGCAAGTCGGCGTTTGTCAAT
GTTAGAGTGCTAGATACGCCAGGCCCTCCACAGAATCTAAAGATAAAGGAGGTCACAAAGACATCTGTCACATTGACGTGGGAGCCTCCTCTCCTGGATGGAGGCTCAAAAATAAAGAACTATATTGTTGAGAAGCGAGAGTCCACAAGAAAGGCATAGCGATCGCACATGGGCTAGCGGA
CTTTGTACACTGGATACTAGTCCCATTACCCTGGTAGATAAGTAGCATGGCGGGTTAATCATTAACTACAAGGAACCCCTAGTGATGGAGTTGGCCACTCCCTCTCTGCGCGCTCGCTCGCTCACTGAGGCCGGGCGACCAAAGGTCGCCCGACGCCCGGGCTTTGCCCGGGCGGCCTCAGTG
AGCGAGCGAGCGCGCAG

**AAV-HDR-mScarlet-PLN-WT** - inserts mScarlet at the N-terminus of wildtype Pln.

CTGCGCGCTCGCTCGCTCACTGAGGCCGCCCGGGCAAAGCCCGGGCGTCGGGCGACCTTTGGTCGCCCGGCCTCAGTGAGCGAGCGAGCGCGCAGAGAGGGAGTGGCCAACTCCATCACTAGGGGTTCCTTGTAGTTAATGATTAACCCGCCATGCTACTTATCTACCAGGGTAATGGGCT
CGAGGAGGGCCTATTTCCCATGATTCCTTCATATTTGCATATACGATACAAGGCTGTTAGAGAGATAATTGGAATTAATTTGACTGTAAACACAAAGATATTAGTACAAAATACGTGACGTAGAAAGTAATAATTTCTTGGGTAGTTTGCAGTTTTAAAATTATGTTTTAAAATGGACTATCATAT
GCTTACCGTAACTTGAAAGTATTTCGATTTCTTGGCTTTATATATCTTGTGGAAAGGACGAAACACCGGGAGACAAGTTTTTCTAAGGTTTTAGAGCTAGAAATAGCAAGTTAAAATAAGGCTAGTCCGTTATCAACTTGAAAAAGTGGCACCGAGTCGGTGCTTTTTTGGTACCATTGTTCCTG
CAGGATACATTGTATTTCCAGCACTCTGGTACAAAAATCAGAAAGATTAAACTTCAAATTAGTCTGTAAGTAAAGAGGCTCTGTGTCTCCAGTAAACACCCAATGAACATATACAGTCATCTTTTTAAACCTGTAGCTAATTTTCTGTTAACACGCTTTATTTTGTTGTATTTACTATCAAAGCTTTG
AACCTTACATATTCAGCCATAGTTTAGAAGTTTTGAAAGAAGGCCAGACACAAACCCACCCCTGGTAGCCAAGATTAGCTGAAAACTGGCACTGTGATAGTTTAAAACGATAGTCCTTACTGTGCCTTCTGAGTTTGGAAGTCTTTAGTACTTAACATCTTCTTAAAGTGTTTAAAATAACATTCA
CCAAGAGGCTGGATCTGGGAGGAGTCCGGAGGATGTATATATGATCAAATACACTGAATGATTCTACAAATCTATAAATCATAAAAGAGAATGGCAAGTGGCTAGAATCTGCTTTTTATTTCATATTTTTCACTTAACTATTTTATTTTGGACTAAAACTAGGATCCAAGATATGTTACTATATGC
ATTTCTTTTTACAAGATTATTTCTGAAACATTAATAGTAAATGTTAAAATCAATTGTGTTTACAGAGAAACAGAGCTTTAAAAAAAGTAACCAAGAAAATAGCCCCAAAGAAGCACCTCATAAATCAAGTACTAGCCGCCAAAGCCGTGGCTCTCCCCGCTCACCCAGGACAGTGAGACGCTCA
CACATGAGAGACAGAAGAGAGGCCTTGGGGACTGGAGAAGGGGATGGGATTTAAAATACAGCTGAGGAACGAGCTGCTGCAAAGCTGTGCATATCTACCCAGAGAACGGGTTTAGGTATGAACCTAATCTGTTGACTATTTTTTACATCCCAGGTCTCCTAAAAGAAGACAGCTCACATTTG
GCTGCCTGTTGTCAACTTTTTATCTTTCTCTTGACAACTTAGAAAAACTTGTCTCCCTACTTTTGCCTTCCTGGCATAgccaccATGGTGAGCAAGGGCGAGGCAGTGATCAAGGAGTTCATGCGGTTCAAGGTGCACATGGAGGGCTCCATGAACGGCCACGAGTTCGAGATCGAGGGCGAGG
GCGAGGGaCGCCCCTACGAGGGCACCCAGACCGCCAAGCTGAAGGTGACCAAGGGTGGCCCCCTGCCCTTCTCCTGGGACATCCTGTCCCCTCAGTTCATGTACGGCTCCAGGGCCTTCACCAAGCACCCCGCCGACATCCCCGACTACTATAAGCAGTCCTTCCCCGAGGGCTTCAAGTGGG
AGCGCGTGATGAACTTCGAGGACGGCGGCGCCGTGACCGTGACCCAGGACACCTCCCTGGAGGACGGCACCCTGATCTACAAGGTGAAGCTCCGCGGCACCAACTTCCCTCCTGACGGCCCCGTAATGCAGAAGAAaACAATGGGCTGGGAAGCGTCCACCGAGCGGTTGTACCCCGAGG
ACGGCGTGCTGAAGGGCGACATTAAGATGGCCCTGCGCCTGAAGGACGGCGGaCGCTACCTGGCGGACTTCAAGACCACCTACAAGGCCAAGAAGCCCGTGCAGATGCCCGGCGCCTACAACGTCGACCGCAAGTTGGACATCACCTCCCACAACGAGGACTACACCGTGGTGGAACAGT
ACGAACGCTCCGAGGGtCGCCACTCCACCGGtGGtATGGACGAGCTGTACAAGATGGAAAAAGTGCAATACCTCACTCGCTCGGCTATCAGGAGAGCCTCCACTATTGAAATGCCTCAGCAAGCACGTCAGAATCTCCAGAACCTATTTATCAATTTCTGCCTCATCTTGATATGTCTGCTGCTG
ATCTGCATCATTGTGATGCTTCTGTGAAGAGCTGCCGCCACTCCAGACCTGCAACATGCCAACTCAGCTTAAAGCCGAGCACTCCGTCATGGGGGAAGCAACGCACCGCTGCCTTCCGGCTGAGAACAGCTTTGTGAGGGTCACGATTAAGAGTGAAACTAATTGTTAGCCAGTGTATTCATC
TGCTGGATCTTGTAAACAAACTTTACAGTGGCTAAGTGTGCATAATGTAGCTGCCAATTTCCTCAACATGGCTCATAAATTTCTTTCCTAAATATTTCTGACAATGAAGTATGAGTTTTAATTTTGAAACACCACTACCAAAAAAGTTCACTTTCTATATCCAACTCTAGCAAGGTCACTTTATAGCT
CCAGTCTGGGGAATAAATGGAATATATACTGCATTGCCACTACCTTTTTTATAGGAACTAAGTGTTCTTCAGTTCTCCACAAAGTAGAAAATAGTCATTTAAATGAAGTGTTCTTATTGAAAATAGCTCACTGGGTCCTCACATAGATGACACAATGATGACGATGATGATGATGACGACGATGA
TGATGATGACGACGATTCAAATCTCTTGGTCACTACTGTCAGAACAAATACCTTGAAAAAATGTTGTGCCCTTTTTCTACACACTAGTATTTTTAAGCTTATGAGAGTTAAAGGGGGAAATGTAAGAACATATTCTCATATAAAATTATTCACTAATTTTGCTCAAGGGCTGTGTACATGTACATG
TACATTAACCACAAATCTGCTCTCTCTAGGGTTTGATCAACAAGCAGGAACATCTTGCTGTGTTGGCTGTGATAATCACTAAAGATCCAGAGTCTCACCTGAAGCTTGTCGCTCTCATGTATCTCCTTGGCGATCGCACATGGGCTAGCGGACTTTGTACACTGGATACTAGTCCCATTACCCTGG
TAGATAAGTAGCATGGCGGGTTAATCATTAACTACAAGGAACCCCTAGTGATGGAGTTGGCCACTCCCTCTCTGCGCGCTCGCTCGCTCACTGAGGCCGGGCGACCAAAGGTCGCCCGACGCCCGGGCTTTGCCCGGGCGGCCTCAGTGAGCGAGCGAGCGCGCAG

**AAV-HDR-mScarlet-PLN-R14Del** - inserts mScarlet at the N-terminus of R14Del Pln.
CTGCGCGCTCGCTCGCTCACTGAGGCCGCCCGGGCAAAGCCCGGGCGTCGGGCGACCTTTGGTCGCCCGGCCTCAGTGAGCGAGCGAGCGCGCAGAGAGGGAGTGGCCAACTCCATCACTAGGGGTTCCTTGTAGTTAATGATTAACCCGCCATGCTACTTATCTACCAGGGTAATGGGCTCGAGGAGGG
CCTATTTCCCATGATTCCTTCATATTTGCATATACGATACAAGGCTGTTAGAGAGATAATTGGAATTAATTTGACTGTAAACACAAAGATATTAGTACAAAATACGTGACGTAGAAAGTAATAATTTCTTGGGTAGTTTGCAGTTTTAAAATTATGTTTTAAAATGGACTATCATATGCTTACCGTAACTTGAAA
GTATTTCGATTTCTTGGCTTTATATATCTTGTGGAAAGGACGAAACACCGGGAGACAAGTTTTTCTAAGGTTTTAGAGCTAGAAATAGCAAGTTAAAATAAGGCTAGTCCGTTATCAACTTGAAAAAGTGGCACCGAGTCGGTGCTTTTTTGGTACCATTGTTCCTGCAGGATACATTGTATTTCCAGCACTCT
GGTACAAAAATCAGAAAGATTAAACTTCAAATTAGTCTGTAAGTAAAGAGGCTCTGTGTCTCCAGTAAACACCCAATGAACATATACAGTCATCTTTTTAAACCTGTAGCTAATTTTCTGTTAACACGCTTTATTTTGTTGTATTTACTATCAAAGCTTTGAACCTTACATATTCAGCCATAGTTTAGAAGTTTTGA
AAGAAGGCCAGACACAAACCCACCCCTGGTAGCCAAGATTAGCTGAAAACTGGCACTGTGATAGTTTAAAACGATAGTCCTTACTGTGCCTTCTGAGTTTGGAAGTCTTTAGTACTTAACATCTTCTTAAAGTGTTTAAAATAACATTCACCAAGAGGCTGGATCTGGGAGGAGTCCGGAGGATGTATATATG
ATCAAATACACTGAATGATTCTACAAATCTATAAATCATAAAAGAGAATGGCAAGTGGCTAGAATCTGCTTTTTATTTCATATTTTCACTTAACTATTTTATTTTGGACTAAAACTAGGATCCAAGATATGTTACTATATGCATTTCTTTTTACAAGATTATTTCTGAAACATTAATAGTAAATGTTAAAATCAAT
TGTGTTTACAGAGAAACAGAGCTTTAAAAAAAGTAACCAAGAAAATAGCCCCAAAGAAGCACCTCATAAATCAAGTACTAGCCGCCAAAGCCGTGGCTCTCCCCGCTCACCCAGGACAGTGAGACGCTCACACATGAGAGACAGAAGAGAGGCCTTGGGGACTGGAGAAGGGGATGGGATTTAAAATAC
AGCTGAGGAACGAGCTGCTGCAAAGCTGTGCATATCTACCCAGAGAACGGGTTTAGGTATGAACCTAATCTGTTGACTATTTTTTACATCCCAGGTCTCCTAAAAGAAGACAGCTCACATTTGGCTGCCTGTTGTCAACTTTTTATCTTTCTCTTGACAACTTAGAAAAACTTGTCTCCCTACTTTTGCCTTCCTG
GCATAgccaccATGGTGAGCAAGGGCGAGGCAGTGATCAAGGAGTTCATGCGGTTCAAGGTGCACATGGAGGGCTCCATGAACGGCCACGAGTTCGAGATCGAGGGCGAGGGCGAGGGaCGCCCCTACGAGGGCACCCAGACCGCCAAGCTGAAGGTGACCAAGGGTGGCCCCCTGCCCTTCTCCTGGG
ACATCCTGTCCCCTCAGTTCATGTACGGCTCCAGGGCCTTCACCAAGCACCCCGCCGACATCCCCGACTACTATAAGCAGTCCTTCCCCGAGGGCTTCAAGTGGGAGCGCGTGATGAACTTCGAGGACGGCGGCGCCGTGACCGTGACCCAGGACACCTCCCTGGAGGACGGCACCCTGATCTACAAGGTGA
AGCTCCGCGGCACCAACTTCCCTCCTGACGGCCCCGTAATGCAGAAGAAaACAATGGGCTGGGAAGCGTCCACCGAGCGGTTGTACCCCGAGGACGGCGTGCTGAAGGGCGACATTAAGATGGCCCTGCGCCTGAAGGACGGCGGaCGCTACCTGGCGGACTTCAAGACCACCTACAAGGCCAAGAAGC
CCGTGCAGATGCCCGGCGCCTACAACGTCGACCGCAAGTTGGACATCACCTCCCACAACGAGGACTACACCGTGGTGGAACAGTACGAACGCTCCGAGGGtCGCCACTCCACCGGtGGtATGGACGAGCTGTACAAGATGGAAAAAGTGCAATACCTCACTCGCTCGGCTATCAGGGCCTCCACTATTGAAA
TGCCTCAGCAAGCACGTCAGAATCTCCAGAACCTATTTATCAATTTCTGCCTCATCTTGATATGTCTGCTGCTGATCTGCATCATTGTGATGCTTCTGTGAAGAGCTGCCGCCACTCCAGACCTGCAACATGCCAACTCAGCTTAAAGCCGAGCACTCCGTCATGGGGGAAGCAACGCACCGCTGCCTTCCGGCT
GAGAACAGCTTTGTGAGGGTCACGATTAAGAGTGAAACTAATTGTTAGCCAGTGTATTCATCTGCTGGATCTTGTAAACAAACTTTACAGTGGCTAAGTGTGCATAATGTAGCTGCCAATTTCCTCAACATGGCTCATAAATTTCTTTCCTAAATATTTCTGACAATGAAGTATGAGTTTTAATTTTGAAACACC
ACTACCAAAAAAGTTCACTTTCTATATCCAACTCTAGCAAGGTCACTTTATAGCTCCAGTCTGGGGAATAAATGGAATATATACTGCATTGCCACTACCTTTTTTATAGGAACTAAGTGTTCTTCAGTTCTCCACAAAGTAGAAAATAGTCATTTAAATGAAGTGTTCTTATTGAAAATAGCTCACTGGGTCCTCA
CATAGATGACACAATGATGACGATGATGATGATGACGACGATGATGATGATGACGACGATTCAAATCTCTTGGTCACTACTGTCAGAACAAATACCTTGAAAAAATGTTGTGCCCTTTTTCTACACACTAGTATTTTTAAGCTTATGAGAGTTAAAGGGGGAAATGTAAGAACATATTCTCATATAAAATTATT
CACTAATTTTGCTCAAGGGCTGTGTACATGTACATGTACATTAACCACAAATCTGCTCTCTCTAGGGTTTGATCAACAAGCAGGAACATCTTGCTGTGTTGGCTGTGATAATCACTAAAGATCCAGAGTCTCACCTGAAGCTTGTCGCTCTCATGTATCTCCTTGGCGATCGCACATGGGCTAGCGGACTTTGTA
CACTGGATACTAGTCCCATTACCCTGGTAGATAAGTAGCATGGCGGGTTAATCATTAACTACAAGGAACCCCTAGTGATGGAGTTGGCCACTCCCTCTCTGCGCGCTCGCTCGCTCACTGAGGCCGGGCGACCAAAGGTCGCCCGACGCCCGGGCTTTGCCCGGGCGGCCTCAGTGAGCGAGCGAGCGCGC
AG

**AAV-HDR-SNAP-PLN-WT** - inserts SNAP-tag at the N-terminus of wildtype Pln.
CTGCGCGCTCGCTCGCTCACTGAGGCCGCCCGGGCAAAGCCCGGGCGTCGGGCGACCTTTGGTCGCCCGGCCTCAGTGAGCGAGCGAGCGCGCAGAGAGGGAGTGGCCAACTCCATCACTAGGGGTTCCTTGTAGTTAATGATTAACCCGCCATGCTACTTATCTACCAGGGTAATGGGCTCGAGGAGGG
CCTATTTCCCATGATTCCTTCATATTTGCATATACGATACAAGGCTGTTAGAGAGATAATTGGAATTAATTTGACTGTAAACACAAAGATATTAGTACAAAATACGTGACGTAGAAAGTAATAATTTCTTGGGTAGTTTGCAGTTTTAAAATTATGTTTTAAAATGGACTATCATATGCTTACCGTAACTTGAAA
GTATTTCGATTTCTTGGCTTTATATATCTTGTGGAAAGGACGAAACACCGGGAGACAAGTTTTTCTAAGGTTTTAGAGCTAGAAATAGCAAGTTAAAATAAGGCTAGTCCGTTATCAACTTGAAAAAGTGGCACCGAGTCGGTGCTTTTTTGGTACCATTGTTCCTGCAGGATACATTGTATTTCCAGCACTCT
GGTACAAAAATCAGAAAGATTAAACTTCAAATTAGTCTGTAAGTAAAGAGGCTCTGTGTCTCCAGTAAACACCCAATGAACATATACAGTCATCTTTTTAAACCTGTAGCTAATTTTCTGTTAACACGCTTTATTTTGTTGTATTTACTATCAAAGCTTTGAACCTTACATATTCAGCCATAGTTTAGAAGTTTTGA
AAGAAGGCCAGACACAAACCCACCCCTGGTAGCCAAGATTAGCTGAAAACTGGCACTGTGATAGTTTAAAACGATAGTCCTTACTGTGCCTTCTGAGTTTGGAAGTCTTTAGTACTTAACATCTTCTTAAAGTGTTTAAAATAACATTCACCAAGAGGCTGGATCTGGGAGGAGTCCGGAGGATGTATATATG
ATCAAATACACTGAATGATTCTACAAATCTATAAATCATAAAAGAGAATGGCAAGTGGCTAGAATCTGCTTTTTATTTCATATTTTCACTTAACTATTTTATTTTGGACTAAAACTAGGATCCAAGATATGTTACTATATGCATTTCTTTTTACAAGATTATTTCTGAAACATTAATAGTAAATGTTAAAATCAAT
TGTGTTTACAGAGAAACAGAGCTTTAAAAAAAGTAACCAAGAAAATAGCCCCAAAGAAGCACCTCATAAATCAAGTACTAGCCGCCAAAGCCGTGGCTCTCCCCGCTCACCCAGGACAGTGAGACGCTCACACATGAGAGACAGAAGAGAGGCCTTGGGGACTGGAGAAGGGGATGGGATTTAAAATAC
AGCTGAGGAACGAGCTGCTGCAAAGCTGTGCATATCTACCCAGAGAACGGGTTTAGGTATGAACCTAATCTGTTGACTATTTTTTACATCCCAGGTCTCCTAAAAGAAGACAGCTCACATTTGGCTGCCTGTTGTCAACTTTTTATCTTTCTCTTGACAACTTAGAAAAACTTGTCTCCCTACTTTTGCCTTCCTG
GCATAgccaccatggacaaagactgcgaaatgaagcgcaccaccctggatagccctctgggcaagctggaactgtctgggtgcgaacagggcctgcaccgtatcatcttcctgggcaaaggaacatctgccgccgacgccgtggaagtgcctgccccagccgccgtgctgggcggaccagagccactgatgcaggccaccgcctggctcaacgcctactttcaccagcctgaggccatcgagg
agttccctgtgccagccctgcaccacccagtgttccagcaggagagctttacccgccaggtgctgtggaaactgctgaaagtggtgaagttcggagaggtcatcagctacagccacctggccgccctggccggcaatcccgccgccaccgccgccgtgaaaaccgccctgagcggaaatcccgtgcccattctgatcccctgccaccgggtggtgcagggcgacctggacgtgggggctacgagg
gcgggctcgccgtgaaagagtggctgctggcccacgagggccacagactgggcaagcctgggctgggtATGGAAAAAGTGCAATACCTCACTCGCTCGGCTATCAGGAGAGCCTCCACTATTGAAATGCCTCAGCAAGCACGTCAGAATCTCCAGAACCTATTTATCAATTTCTGCCTCATCTTGATATGTCTGCTGCTGATCTGCAT
CATTGTGATGCTTCTGTGAAGAGCTGCCGCCACTCCAGACCTGCAACATGCCAACTCAGCTTAAAGCCGAGCACTCCGTCATGGGGGAAGCAACGCACCGCTGCCTTCCGGCTGAGAACAGCTTTGTGAGGGTCACGATTAAGAGTGAAACTAATTGTTAGCCAGTGTATTCATCTGCTGGATCTTGTAAACA
AACTTTACAGTGGCTAAGTGTGCATAATGTAGCTGCCAATTTCCTCAACATGGCTCATAAATTTCTTTCCTAAATATTTCTGACAATGAAGTATGAGTTTTAATTTTGAAACACCACTACCAAAAAAGTTCACTTTCTATATCCAACTCTAGCAAGGTCACTTTATAGCTCCAGTCTGGGGAATAAATGGAATATA
TACTGCATTGCCACTACCTTTTTTATAGGAACTAAGTGTTCTTCAGTTCTCCACAAAGTAGAAAATAGTCATTTAAATGAAGTGTTCTTATTGAAAATAGCTCACTGGGTCCTCACATAGATGACACAATGATGACGATGATGATGATGACGACGATGATGATGATGACGACGATTCAAATCTCTTGGTCACTA
CTGTCAGAACAAATACCTTGAAAAAATGTTGTGCCCTTTTTCTACACACTAGTATTTTTAAGCTTATGAGAGTTAAAGGGGGAAATGTAAGAACATATTCTCATATAAAATTATTCACTAATTTTGCTCAAGGGCTGTGTACATGTACATGTACATTAACCACAAATCTGCTCTCTCTAGGGTTTGATCAACAAG
CAGGAACATCTTGCTGTGTTGGCTGTGATAATCACTAAAGATCCAGAGTCTCACCTGAAGCTTGTCGCTCTCATGTATCTCCTTGGCGATCGCACATGGGCTAGCGGACTTTGTACACTGGATACTAGTCCCATTACCCTGGTAGATAAGTAGCATGGCGGGTTAATCATTAACTACAAGGAACCCCTAGTGAT
GGAGTTGGCCACTCCCTCTCTGCGCGCTCGCTCGCTCACTGAGGCCGGGCGACCAAAGGTCGCCCGACGCCCGGGCTTTGCCCGGGCGGCCTCAGTGAGCGAGCGAGCGCGCAG

**AAV-HDR-SNAP-PLN-R14Del** - inserts SNAP-tag at the N-terminus of R14Del Pln.
CTGCGCGCTCGCTCGCTCACTGAGGCCGCCCGGGCAAAGCCCGGGCGTCGGGCGACCTTTGGTCGCCCGGCCTCAGTGAGCGAGCGAGCGCGCAGAGAGGGAGTGGCCAACTCCATCACTAGGGGTTCCTTGTAGTTAATGATTAACCCGCCATGCTACTTATCTACCAGGGTAATGGGCTCGAGGAGGG
CCTATTTCCCATGATTCCTTCATATTTGCATATACGATACAAGGCTGTTAGAGAGATAATTGGAATTAATTTGACTGTAAACACAAAGATATTAGTACAAAATACGTGACGTAGAAAGTAATAATTTCTTGGGTAGTTTGCAGTTTTAAAATTATGTTTTAAAATGGACTATCATATGCTTACCGTAACTTGAAA
GTATTTCGATTTCTTGGCTTTATATATCTTGTGGAAAGGACGAAACACCGGGAGACAAGTTTTTCTAAGGTTTTAGAGCTAGAAATAGCAAGTTAAAATAAGGCTAGTCCGTTATCAACTTGAAAAAGTGGCACCGAGTCGGTGCTTTTTTGGTACCATTGTTCCTGCAGGATACATTGTATTTCCAGCACTCT
GGTACAAAAATCAGAAAGATTAAACTTCAAATTAGTCTGTAAGTAAAGAGGCTCTGTGTCTCCAGTAAACACCCAATGAACATATACAGTCATCTTTTTAAACCTGTAGCTAATTTTCTGTTAACACGCTTTATTTTGTTGTATTTACTATCAAAGCTTTGAACCTTACATATTCAGCCATAGTTTAGAAGTTTTGA
AAGAAGGCCAGACACAAACCCACCCCTGGTAGCCAAGATTAGCTGAAAACTGGCACTGTGATAGTTTAAAACGATAGTCCTTACTGTGCCTTCTGAGTTTGGAAGTCTTTAGTACTTAACATCTTCTTAAAGTGTTTAAAATAACATTCACCAAGAGGCTGGATCTGGGAGGAGTCCGGAGGATGTATATATG
ATCAAATACACTGAATGATTCTACAAATCTATAAATCATAAAAGAGAATGGCAAGTGGCTAGAATCTGCTTTTTATTTCATATTTTCACTTAACTATTTTATTTTGGACTAAAACTAGGATCCAAGATATGTTACTATATGCATTTCTTTTTACAAGATTATTTCTGAAACATTAATAGTAAATGTTAAAATCAAT
TGTGTTTACAGAGAAACAGAGCTTTAAAAAAAGTAACCAAGAAAATAGCCCCAAAGAAGCACCTCATAAATCAAGTACTAGCCGCCAAAGCCGTGGCTCTCCCCGCTCACCCAGGACAGTGAGACGCTCACACATGAGAGACAGAAGAGAGGCCTTGGGGACTGGAGAAGGGGATGGGATTTAAAATAC
AGCTGAGGAACGAGCTGCTGCAAAGCTGTGCATATCTACCCAGAGAACGGGTTTAGGTATGAACCTAATCTGTTGACTATTTTTTACATCCCAGGTCTCCTAAAAGAAGACAGCTCACATTTGGCTGCCTGTTGTCAACTTTTTATCTTTCTCTTGACAACTTAGAAAAACTTGTCTCCCTACTTTTGCCTTCCTG
GCATAgccaccatggacaaagactgcgaaatgaagcgcaccaccctggatagccctctgggcaagctggaactgtctgggtgcgaacagggcctgcaccgtatcatcttcctgggcaaaggaacatctgccgccgacgccgtggaagtgcctgccccagccgccgtgctgggcggaccagagccactgatgcaggccaccgcctggctcaacgcctactttcaccagcctgaggccatcgagg
agttccctgtgccagccctgcaccacccagtgttccagcaggagagctttacccgccaggtgctgtggaaactgctgaaagtggtgaagttcggagaggtcatcagctacagccacctggccgccctggccggcaatcccgccgccaccgccgccgtgaaaaccgccctgagcggaaatcccgtgcccattctgatcccctgccaccgggtggtgcagggcgacctggacgtgggggctacgagg
gcgggctcgccgtgaaagagtggctgctggcccacgagggccacagactgggcaagcctgggctgggtATGGAAAAAGTGCAATACCTCACTCGCTCGGCTATCAGGGCCTCCACTATTGAAATGCCTCAGCAAGCACGTCAGAATCTCCAGAACCTATTTATCAATTTCTGCCTCATCTTGATATGTCTGCTGCTGATCTGCATCATT
GTGATGCTTCTGTGAAGAGCTGCCGCCACTCCAGACCTGCAACATGCCAACTCAGCTTAAAGCCGAGCACTCCGTCATGGGGGAAGCAACGCACCGCTGCCTTCCGGCTGAGAACAGCTTTGTGAGGGTCACGATTAAGAGTGAAACTAATTGTTAGCCAGTGTATTCATCTGCTGGATCTTGTAAACAAAC
TTTACAGTGGCTAAGTGTGCATAATGTAGCTGCCAATTTCCTCAACATGGCTCATAAATTTCTTTCCTAAATATTTCTGACAATGAAGTATGAGTTTTAATTTTGAAACACCACTACCAAAAAAGTTCACTTTCTATATCCAACTCTAGCAAGGTCACTTTATAGCTCCAGTCTGGGGAATAAATGGAATATATAC
TGCATTGCCACTACCTTTTTTATAGGAACTAAGTGTTCTTCAGTTCTCCACAAAGTAGAAAATAGTCATTTAAATGAAGTGTTCTTATTGAAAATAGCTCACTGGGTCCTCACATAGATGACACAATGATGACGATGATGATGATGACGACGATGATGATGATGACGACGATTCAAATCTCTTGGTCACTACTG
TCAGAACAAATACCTTGAAAAAATGTTGTGCCCTTTTTCTACACACTAGTATTTTTAAGCTTATGAGAGTTAAAGGGGGAAATGTAAGAACATATTCTCATATAAAATTATTCACTAATTTTGCTCAAGGGCTGTGTACATGTACATGTACATTAACCACAAATCTGCTCTCTCTAGGGTTTGATCAACAAGCAG
GAACATCTTGCTGTGTTGGCTGTGATAATCACTAAAGATCCAGAGTCTCACCTGAAGCTTGTCGCTCTCATGTATCTCCTTGGCGATCGCACATGGGCTAGCGGACTTTGTACACTGGATACTAGTCCCATTACCCTGGTAGATAAGTAGCATGGCGGGTTAATCATTAACTACAAGGAACCCCTAGTGATGGA
GTTGGCCACTCCCTCTCTGCGCGCTCGCTCGCTCACTGAGGCCGGGCGACCAAAGGTCGCCCGACGCCCGGGCTTTGCCCGGGCGGCCTCAGTGAGCGAGCGAGCGCGCAG

**AAV-HDR-Tnni1-2xHA-Enh** - inserts 2xHA-tag into Tnni1 C-terminus, and enhancer pool into Tnni1 3' UTR. N = enhancer and barcode.
CTGCGCGCTCGCTCGCTCACTGAGGCCGCCCGGGCAAAGCCCGGGCGTCGGGCGACCTTTGGTCGCCCGGCCTCAGTGAGCGAGCGAGCGCGCAGAGAGGGAGTGGCCAACTCCATCACTAGGGGTTCCTTGTAGTTAATGATTAACCCGCCATGCTACTTATCTACCAGGGTAATGGGCTCGAGGAGGG
CCTATTTCCCATGATTCCTTCATATTTGCATATACGATACAAGGCTGTTAGAGAGATAATTGGAATTAATTTGACTGTAAACACAAAGATATTAGTACAAAATACGTGACGTAGAAAGTAATAATTTCTTGGGTAGTTTGCAGTTTTAAAATTATGTTTTAAAATGGACTATCATATGCTTACCGTAACTTGAAA
GTATTTCGATTTCTTGGCTTTATATATCTTGTGGAAAGGACGAAACACCCTCTAAGCACCTCTACTGGGGTTTTAGAGCTAGAAATAGCAAGTTAAAATAAGGCTAGTCCGTTATCAACTTGAAAAAGTGGCACCGAGTCGGTGCTTTTTTGGTACCATTGTTCCTGCAGGGCCCTACTGGGCTCTAAGCACAA
GGTGTCCATGGATCTGCGGGCCAACCTCAAGTCTGTGAAGAAAGAAGACACAGAAAAGGTGAGGGCCCCTCTCCTCAGACCCGCCTACTGGCCTCTATCTGCCACTTCTTACATTGCCTGAGGCGCTCTAGTCAGTGTGTGAATCAGGATATTCTAGGATCCCCTAGGAGGCCAGTGTCAGAAGAGAGACTA
CTGGCTGCTTCCAGTCTGAGGGGACAATGGGATGAGAGAAAGGCTGACTGGGAAATTAAAACTTGCTTCCCATATCAGTTAAGGGGTAACCATTACCCCTGTCACTTCAGCTTCTCGTTGTGGCTATAGGAACATGAGTGCATAGAAATTGGAGAACCGAGTTCAATTAACTTGCCTAGCAAACAACTACTTC
AAATTTGTTGGTCTCACAGAACATGTTTAGAGCTGACAGGCTGTGAGGAGCATGGTGAGAGAGCAATTTATCTGGCCCAGAGAGCTTAATAAAGACAAACAGGCCAAGCCAGTATGCTGACAAAATTGAAAACAAGTTAAATGAATCAGGAGAGGCTTCTTGGAGGAGGTGACCTTGAACCAACTGCAGG
GTGTCCCACAGGAGGGTGTTGTGGATCCTGGAGGAGAGGTTGGCCTGTCATATACTGGGAACCAGTTTTGTGACCCTGTGACAGTTACCTAGACCCTCCCATCCACACAGCCTCATTCTGTGCTCCTGTCTGGCCTATTCATCTTGTATCATGTGCTGGGCAGGGCCATCAGGATTTTTTAGCTCTCTGAGGAG
GTAGATTTTCTCCCATCTACTTTCTCTTGAGCTTGTGGGTCCTTTGTTGCCCCTGCGGCCTGGTGAGGCCCCTGCAGTGCTGACATGTTTGTGGACTGCCCACAGGAGCGGCCAGTCGAGGTAGGAGACTGGAGGAAGAATGTGGAGGCTATGTCTGGCATGGAAGGCCGCAAGAAGATGTTCGACGCTGCT
AAGTCCCCGACCTCCCAGTACCCCTACGACGTGCCCGACTACGCTTACCCTTACGATGTGCCTGATTATGCCTAGGCGGCCGCNNNNNNNNNNNNNNNNNNNNNNNNNNNNNNNNNNNNNNNNNNNNNNNNNNNNNNNNNNNNNNNNNNNNNNNNNNNNNNNNNNNNNNNNNNN
NNNNNNNNNNNNNNNNNNNNNNNNNNNNNNNNNNNNNNNNNNNNNNNNNNNNNNNNNNNNNNNNNNNNNNNNNNNNNNNNNNNNNNNNNNNNNNNNNNNNNNNNNNNNNNNNNNNNNNNNNNNNNNNNNNNNNNNNNNN
NNNNNNNNNNNNNNNNNNNNNNNNNNNNNNNNNNNNNNNNNNNNNNNNNNNNNNNNNNNNNNNNNNNNNNNNNNNNNNNNNNNNNNNNNNNNNNNNNNNNNNNNNNNNATCGATAATCGGCCGGCCGTCAGCGATCGCNNNN
NNNNNNGGCGCGCCCAGGTGAGTAGTAGAGGTGCTTAGAGTGTTCTTGGggtccctcagagtaccccagggtactcaggagaagggatgggaggcagatccaatgtgcagaactccagcctgctccagtaccttcagacactgggcacagcatgtctatgccagatctttttgtccccttatcagtaaaacttgggcttgtatgtgatcagtggtcctggaactggcagcat
ctatgtgttcctggttgcaaatgactgaactggaaactggggggaccctagaggaccctgagacaggcaagagtctgagagtacagggttaggtgatgcacaaggtccctccagctctaacgcttttctaactcagtcttttctttcacttgtcacttggtgccatacttggcttccataccctcaaagaagctcccccagtcactactcccagagaaacagcagttaagagatgttcaaacccccccc
ccactccagagaaatagctgcaggaactgctgaagcagtggaattgggccataaaagccctgctgtgggccctaggaggggaggtgtaatagccaggtctacaggccacacagagccaacttcttcacagagcagatgttagctagtgataggtgagggttcacacatgaaggggaagggctgagggtgcaaaggtcagctatgggagggaaccggtctggaagatggccaggacaggttgg
agggaaggagctagagagggaggcagcttgggatctggacagagcagctaagtcccatgggtggtacaagggagaagccccagggactatgggggctgtgtgtgctgtgtgttaatcagggttacctaaacagcacgcatcactgggatctatctgatggcatctcctcttttgtcctgacaggttggcttgctgtatgacactctgtgtcatgctttggtccctttccagccccactcacctatctttct
gctcgtgcctggaacatccctaatatgcGCTCCCTCCCTCCAGCCTATGCGATCGCACATGGGCTAGCGGACTTTGTACACTGGATACTAGTCCCATTACCCTGGTAGATAAGTAGCATGGCGGGTTAATCATTAACTACAAGGAACCCCTAGTGATGGAGTTGGCCACTCCCTCTCTGCGCGCTCGCTCGCTCACTGAG
GCCGGGCGACCAAAGGTCGCCCGACGCCCGGGCTTTGCCCGGGCGGCCTCAGTGAGCGAGCGAGCGCGCAG

**AAV-Enh-Prom-mCherry-bc-polyA** - MPRA vector carrying barcoded enhancer:promoter pairs, driving expression of mCherry reporter transcript. N = enhancer, promoter, promoter barcode, enhancer barcode respectively.
ctgcgcgctcgctcgctcactgaggccgcccgggcaaagcccgggcggcctcagtgagcgagcgagcgcgcagagagggagtggccaactccatcactaggggttccttgtagttaatgattaacccgccatgctacttatctaccagggtaatggggatcctctagaactatagctagaattcgtttggtggcactggagcggccgcNNNNNNNNNNNNNNNNNNNNNNNNNN
NNNNNNNNNNNNNNNNNNNNNNNNNNNNNNNNNNNNNNNNNNNNNNNNNNNNNNNNNNNNNNNNNNNNNNNNNNNNNNNNNNNNNNNNNNNNNNNNNNNNNNNNNNNNNNNNNNNNNNNNNNNNNNNNNNNNNNNNNNN
NNNNNNNNNNNNNNNNNNNNNNNNNNNNNNNNNNNNNNNNNNNNNNNNNNNNNNNNNNNNNNNNNNNNNNNNNNNNNNNNNNNNNNNNNNNNNNNNNNNNNNNNNNNNNNNNNNNNNNNNNNNNNNNNNNNNNNNNNNN
NNNNNNNNNNNNNNNNNNNNNNNNNNNNNNNNNNNNNNNNNNNNNatcgatNNNNNNNNNNNNNNNNNNNNNNNNNNNNNNNNNNNNNNNNNNNNNNNNNNNNNNNNNNNNNNNNNNNNNNNNNNNNNNNNNNNNNNNNNNNNNN
NNNNNNNNNNNNNNNNNNNNNNNNNNNNNNNNNNNNNNNNNNNNNNNNNNNNNNNNNNNNNNNNNNNNNNNNNNNNNNNNNNNNNNNgccaccatggtgagcaagggcgaggaggataacatggccatcatcaaggagttcatgcgcttcaaggtgcacatggagggctccgtgaacggccacgagtttgagattgaaggcgaaggcgagggcagac
cctacgagggaacacagaccgccaagctgaaggtgaccaagggtggcccccctgcccttcgcctgggacatcctgtcccctcagttcatgtacggctccaaggcctacgtgaagcaccccgccgacatccccgactacttgaagctgtccttccccgagggcttcaagtgggagcgcgtgatgaacttcgaggacggcggcgtggtgaccgtgacccaggactcctccctgcaggacggcgagttca
tctacaaggtgaagctgcgcggcaccaacttcccctccgacggccccgtaatgcagaagaagacaatgggctgggaggcctcctccgagcggatgtaccccgaggacggcgccctgaagggcgagatcaagcagaggctgaagctgaaggacggcggccactacgacgctgaggtcaagaccacctacaaggccaagaagcccgtgcagctgcccggcgcctacaacgtcaacatcaagttg
gacatcacctcccacaacgaggactacaccatcgtggaacagtacgaacgcgccgagggtcgacactccaccggcggcatggacgagctgtacaagtagNNNNNNNNNNgcgatcgcNNNNNNNNNNggcgcgcctagccatctcgaggacggggtgaactacgcctgaggatccgatctttttccctctgccaaaaattatggggacatcatgaagccccttgagcatctgactt
ctggctaataaaggaaatttattttcattgcaatagtgtgttggaattttttgtgtctctcactcggaagcaattcgttgatctgaatttcgaccacccataatacccattaccctggtagataagtagcatggcgggttaatcattaactacaaggaacccctagtgatggagttggccactccctctctgcgcgctcgctcgctcactgaggccgggcgaccaaaggtcgcccgacgcccgggctttgcc
cgggcggcctcagtgagcgagcgagcgcgcag
